# *PRNP* mutations initially generate an alternatively misfolded PrP species that dissuades prion replication

**DOI:** 10.64898/2026.09.22.753491

**Authors:** Genki Amano, Hamza Arshad, Surabhi Mehra, Matthew E.C. Bourkas, Erica Stuart, Gerold Schmitt-Ulms, Surachai Supattapone, Joel C. Watts

**Author notes:** To whom correspondence should be addressed at: Krembil Discovery Tower, Rm. 4KD481, 60 Leonard Ave., Toronto, ON, Canada, M5T 0S8.

## Abstract

Mutations in the *PRNP* gene, which encodes the prion protein (PrP), cause genetic prion disease. However, how *PRNP* mutations lead to spontaneous prion formation remains poorly understood. Building off the observation that expression of mutant bank vole PrP (BVPrP) in mice causes spontaneous prion disease, we sought to identify early events in prion formation by expressing mutant BVPrP in cultured cells lacking endogenous PrP. Expression of D178N- and E200K-mutant, but not wild-type BVPrP, in CAD5-PrP^-/-^ and N2a-PrP^-/-^ cells generates an <u>a</u>lternatively <u>m</u>isfolded PrP species, which we term PrP^AM^, that is detergent-insoluble and resistant to digestion with the protease thermolysin. PrP^AM^ is also present in the brains of young, asymptomatic knock-in mice expressing mutant BVPrP. However, PrP^AM^ does not appear to be a direct precursor of prions since 1) the prion disease-protective G127V substitution enhances PrP^AM^ levels when coupled with the D178N or E200K mutations; 2) small molecule anti-prion drugs fail to reduce PrP^AM^ accumulation in cells; 3) unlike cells expressing wild-type BVPrP, cells expressing D178N- or E200K-mutant BVPrP are largely resistant to infection with several prion strains; and 4) PrP^AM^-containing cell homogenate does not contain seeds that induce misfolding of wild-type BVPrP in either cells or the RT-QuIC assay. These results raise the possibility that prion disease-causing *PRNP* mutations initially generate a protective misfolded PrP species that is refractory to prion formation, which provides a potential explanation for why genetic prion diseases manifest later in life despite the presence of a mutation from birth.

## Introduction

Prion diseases are fatal neurodegenerative disorders affecting both humans and animals that can manifest either sporadically, genetically, or via an infection. Prions are misfolded conformers of the prion protein (PrP) that induce profound neuropathological changes including spongiform degeneration, astrocytic gliosis, and neuronal loss (1, 2). The cellular (normal) version of PrP (PrP^C^) is a predominantly α-helical glycoprotein that is tethered to the surface of neurons and other brain cell types via a glycophosphatidylinositol (GPI) anchor (3, 4). During prion disease, PrP^C^ misfolds into PrP^Sc^, which is composed of β-sheet rich aggregates that are neurotoxic, infectious, and partially resistant to digestion with proteases such as proteinase K (PK) (5–8). Different prion strains are recognized, each of which represents a distinct structure of PrP^Sc^ aggregates (9, 10). Through a poorly understood process referred to as prion replication, PrP^Sc^ can act as a seed or template to induce the misfolding of PrP^C^ into additional PrP^Sc^ (11). Thus, PrP^Sc^ aggregates are self-propagating, allowing prions to spread from cell-to-cell via a cascade of protein misfolding. Since PrP^C^ is the substrate for prion replication, PrP^C^ expression is absolutely required for prion disease. Accordingly, mice lacking PrP^C^ (PrP^-/-^) are resistant to prion infection (12). There are currently no approved therapies for any of the human or animal prion diseases, although several strategies based on reduction of PrP^C^ levels in the brain are in various stages of development (13–17).

In sporadic human prion disorders such as sporadic Creutzfeldt-Jakob disease (CJD), which constitute ∼90% of cases in humans, the initiating event in the disease process is thought to be the spontaneous misfolding of PrP^C^ to generate a PrP^Sc^ seed that can then propagate (18). Sporadic CJD has an incidence of 1-2 new cases per million population per year, which suggests that spontaneous generation of a stable PrP^Sc^ seed is a rare event and/or that such seeds are normally cleared by proteostatic machinery. In the infectious prion diseases, which constitute ∼1% of human prion disease cases, an exogenous source of PrP^Sc^ initiates the disease process by converting host-expressed PrP^C^ into PrP^Sc^, bypassing the need for seed formation (19). In genetic prion diseases such as genetic CJD, Gerstmann-Sträussler Scheinker syndrome (GSS), and fatal familial insomnia (FFI), which constitute ∼10% of human prion disease cases, mutations are present in the *PRNP* gene, which encodes PrP (20). Mutant PrP is believed to be more prone to spontaneously forming PrP^Sc^ than wild-type (WT) PrP, although precise molecular details of how specific mutations promote PrP^Sc^ formation remain lacking since prion disease-causing mutations do not uniformly alter the structure or stability of PrP^C^ (21–24).

The study of prion disease and the identification of anti-prion therapeutic strategies have been facilitated by the existence of mouse and cellular paradigms that can become infected with prions upon exposure to a pre-existing source of PrP^Sc^ (25, 26). However, roughly 99% of human prion disease cases do not arise via an infection. Thus, models that recapitulate spontaneous prion formation may be more informative when developing therapeutics for the sporadic and genetic prion diseases (27). Two of the most common *PRNP* mutations are D178N and E200K, which can cause either genetic CJD (D178N and E200K) or FFI (D178N) (28–33). Cells expressing D178N- or E200K-mutant PrP do not spontaneously develop PK-resistant PrP species like those found in prion disease brains but do exhibit misfolded PrP species that are insoluble and partially resistant to digestion with low concentrations of PK (34–41). Similarly, transgenic or knock-in mice expressing D178N- or E200K-mutant PrP recapitulate some, but not all aspects of the corresponding genetic human prion disorder (42–48). However, in other cellular paradigms, it has been reported that the E200K mutation does not appreciably alter the biochemical properties of PrP (49–51).

While previous studies have used either mouse or human PrP as the backbone for studying the pathogenic effects of the D178N and E200K *PRNP* mutations, it has become apparent that the bank vole prion protein (BVPrP) may be a better substrate for studying spontaneous prion misfolding. Bank voles and transgenic mice expressing BVPrP can replicate many different prion strains from several different species, leading to the suggestion that BVPrP may be a ‘universal prion acceptor’ (52–59). Moreover, recombinant BVPrP readily polymerizes into aggregates, either spontaneously or upon seeding with PrP^Sc^, and transgenic mice over-expressing WT BVPrP develop spontaneous prion disease (60–66). Transgenic and knock-in mice expressing D178N- or E200K-mutant BVPrP develop spontaneous neurological illness and recapitulate several aspects of authentic prion disease including the presence of PK-resistant PrP species in the brain, spongiform degeneration, and the generation of prion infectivity and seeding activity (67, 68). Thus, we hypothesized that D178N- and E200K-mutant BVPrP may facilitate our understanding of how prions form spontaneously. In this study, we expressed mutant BVPrP in cultured cells lacking endogenous PrP and then assessed their biochemical properties. Our findings suggest that *PRNP* mutations initially generate an alternatively misfolded PrP species, which we term PrP^AM^, that dissuades PrP^Sc^ formation and prion replication.

## Results

### The D178N and E200K mutations induce formation of PrP^AM^, a thermolysin-resistant species

To assess potential early misfolding events caused by the prion disease-causing D178N and E200K mutations, we inserted the mutations into BVPrP (**Fig. 1a**). For all experiments, the I109 polymorphic variant of BVPrP was used because it is more prone to spontaneous misfolding than the M109 variant (60). Murine CAD5 (<u>C</u>ath.<u>a</u>-<u>d</u>ifferentiated) cells, which are a derivative of the Cath.a catecholaminergic neuron-like line (69), are susceptible to a wide range of mouse prion strains (70, 71). Moreover, engineered CAD5-PrP^-/-^ cells expressing PrPs from other species can propagate non-mouse prion strains (63, 72–77). Thus, we speculated that this cell line would be optimal for assessing spontaneous BVPrP misfolding. Wild-type (WT) and mutant BVPrPs were stably expressed in CAD5-PrP^-/-^ cells lacking endogenous expression of mouse PrP (MoPrP). Initially, we generated polyclonal lines containing a pool of stably transfected cells. In line with what has previously been observed in stably transfected N2a neuroblastoma cells lacking PrP expression (N2a-PrP^-/-^) and in the brains of BVPrP(I109) knock-in mice (67, 68), steady-state levels of D178N- and E200K-mutant BVPrP(I109) were lower than those for WT BVPrP(I109) (**Fig. 1b, c**). Similar results were obtained with several different anti-PrP antibodies using both reduced and non-reduced samples (**Fig. S1**). Consistent with previous observations (78), PrP signals were higher in non-reduced samples than reduced samples when using the POM1 antibody that recognizes the α-helical domain in PrP. As in BVPrP(I109) knock-in mice (68), D178N-mutant BVPrP(I109) expressed in CAD5-PrP^-/-^ cells was almost exclusively diglycosylated whereas monoglycosylated species were also present for WT and E200K-mutant BVPrP(I109) (**Fig. 1b**). In cells expressing WT BVPrP(I109), ∼2% of PrP species are detergent-insoluble (**Fig. S2**). However, when normalized to levels of total PrP, relative levels of detergent-insoluble PrP were ∼8-fold higher in cells expressing D178N-mutant BVPrP(I109) and ∼3-4 fold higher in cells expressing E200K-mutant BVPrP(I109) compared to cells expressing WT BVPrP(I109) (**Fig. 1d, e**).

**Figure 1.**
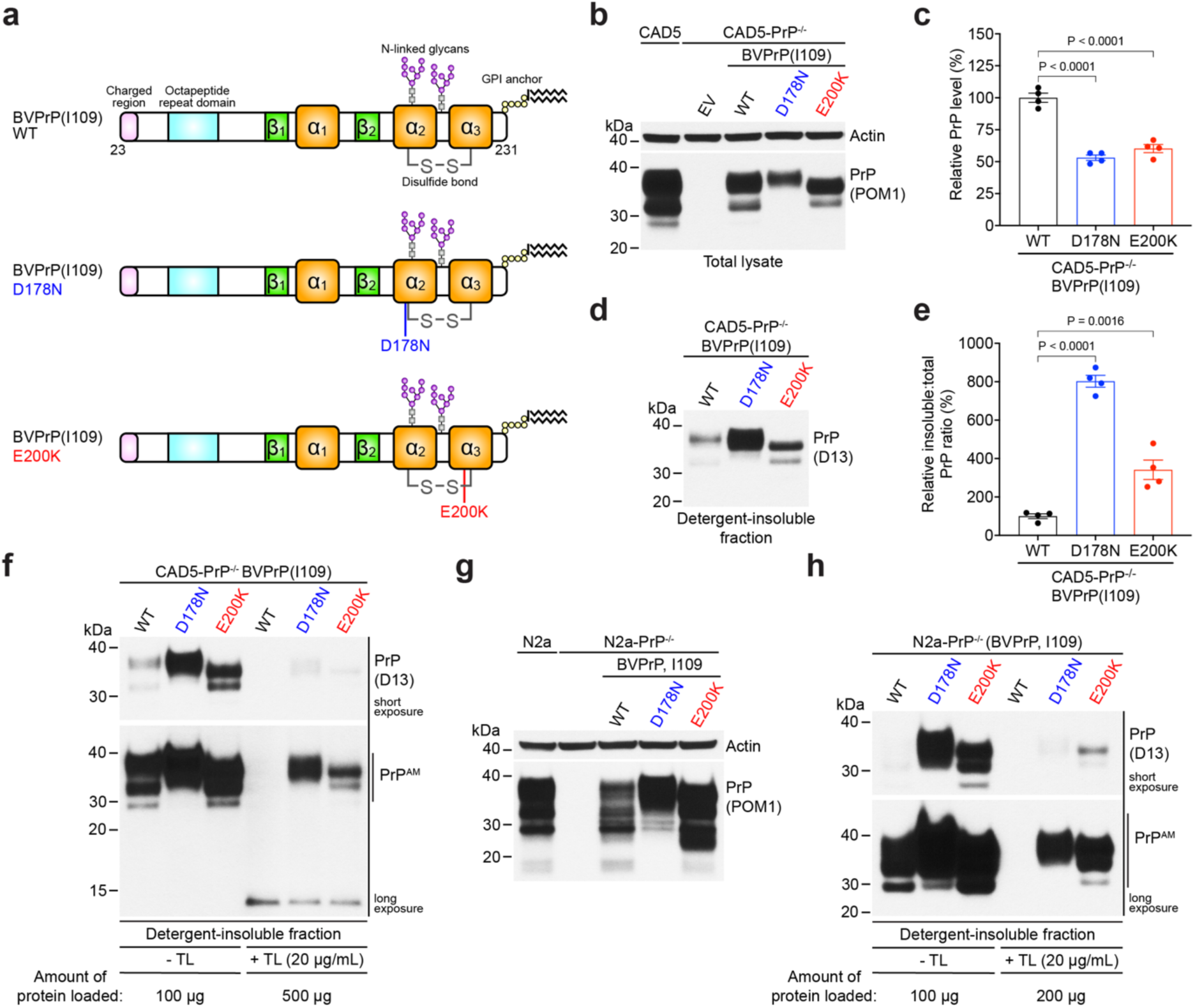
Stably transfected PrP^-/-^ cells expressing mutant bank vole PrP produce a thermolysin-resistant PrP species termed PrP^AM^. **a**) Schematic structures of WT, D178N, and E200K BVPrP(I109). **b**) Representative immunoblot of total PrP levels in lysates from polyclonal lines of CAD5-PrP^-/-^ cells stably transfected with either empty vector (EV) or with the indicated BVPrP(I109) constructs. Lysates from untransfected CAD5 cells are included as a control. The blot was reprobed with an antibody against actin. **c**) Quantification of total PrP levels in lysates from stably transfected CAD5-PrP^-/-^ cells. **d**) Representative immunoblot of detergent-insoluble PrP levels in lysates from CAD5-PrP^-/-^ cells stably transfected with the indicated BVPrP(I109) constructs. **e**) Quantification of detergent-insoluble PrP levels (normalized to total PrP levels) in lysates from stably transfected CAD5-PrP^-/-^ cells. **f**) Representative immunoblot of detergent-insoluble PrP species in lysates from stably transfected CAD5-PrP^-/-^ cells that were either treated (+ TL) or not treated (- TL) with 20 µg/mL thermolysin. Note that 5X more protein was loaded for the TL-digested samples compared to the undigested samples. A detergent-insoluble and TL-resistant PrP species (“PrP^AM^”) is selectively present in cells expressing D178N- or E200K-mutant BVPrP(I109). **g**) Representative immunoblot of total PrP levels in lysates from monoclonal lines of N2a-PrP^-/-^ cells stably transfected with the indicated BVPrP(I109) constructs. Lysates from untransfected N2a and N2a-PrP^-/-^ cells are included as a control. The blot was reprobed with an antibody against actin. **h**) Representative immunoblot of detergent-insoluble PrP species in lysates from stably transfected N2a-PrP^-/-^ cells that were either treated (+ TL) or not treated (- TL) with 20 µg/mL thermolysin.

Both classical and atypical forms of PrP^Sc^ are resistant to digestion with the protease thermolysin (TL) when used at a concentration of 100 µg/mL (79, 80). Thus, we wondered whether the detergent-insoluble PrP species in cells expressing mutant BVPrP(I109) might be resistant to TL digestion. Lysates from cells expressing either WT or E200K-mutant BVPrP(I109) were treated with various concentrations of TL and then the detergent-insoluble fraction was isolated by ultracentrifugation. Whereas no TL-resistant PrP was observed when a TL concentration of 50 µg/mL was used, higher levels of detergent-insoluble PrP species were observed in lysates from cells expressing E200K-mutant BVPrP(I109) compared to cells expressing WT BVPrP(I109) when TL concentrations of 5-25 µg/mL were used (**Fig. S3**). Thus, we decided to use a TL concentration of 20 µg/mL for all subsequent experiments. Cells expressing either D178N- or E200K-mutant BVPrP(I109) selectively exhibited detergent-insoluble PrP species that were resistant to digestion with TL (**Fig. 1f**). This suggests that a subset of D178N-mutant and E200K-mutant BVPrP(I109) spontaneously misfolds into a structure that is detergent-insoluble and resistant to mild TL digestion when expressed in CAD5-PrP^-/-^ cells. We term this species “PrP^AM^” (<u>a</u>lternatively <u>m</u>isfolded PrP) to differentiate it from PrP^Sc^. The molecular weights of detergent-insoluble D178N- and E200K-mutant BVPrP(I109) species were not appreciably altered upon TL digestion, suggesting that PrP^AM^ is composed of full-length, N-glycosylated PrP. This suggests that PrP^AM^ is not simply mutant PrP that has failed quality control mechanisms within the secretory pathway during protein folding, although mutant PrPs are known to be delayed in their exit from the endoplasmic reticulum (41). PrP^AM^ was also observed in lysates from N2a-PrP^-/-^ cells stably expressing either D178N- or E200K-mutant BVPrP(I109), demonstrating that PrP^AM^ production is not restricted to the CAD5 cell line (**Fig. 1g, h**). No PK-resistant PrP species were observed in the CAD5-PrP^-/-^ cell lines expressing D178N-mutant or E200K-mutant BVPrP(I109), even when low concentrations of PK were used (**Fig. S4**), implying that PrP^AM^ is distinct from the PrP^Sc^ aggregates found in prion-infected cells and animals.

To determine whether production of PrP^AM^ is specific to BVPrP, we generated polyclonal stable CAD5-PrP^-/-^ cell lines expressing either WT MoPrP or the MoPrP equivalents of the D178N or E200K mutations (D177N and E199K, respectively). As with the BVPrP(I109) lines, cells expressing D177N-mutant or E199K-mutant MoPrP displayed lower levels of PrP expression than cells expressing WT MoPrP (**Fig. 2a, b**). Likewise, when normalized to levels of total PrP expression, relative levels of detergent-insoluble PrP species were elevated ∼3-fold in cells expressing D177N-mutant MoPrP and ∼2-fold in cells expressing E199K-mutant MoPrP (**Fig. 2c, d**). Consistent with what was observed with the BVPrP-expressing lines, detergent-insoluble and TL-resistant PrP^AM^ species were also present in cells expressing either D177N- or E199K-mutant MoPrP (**Fig. 2e**). Thus, the D178N and E200K mutations specify formation of PrP^AM^ when inserted into PrPs from two different species.

**Figure 2.**
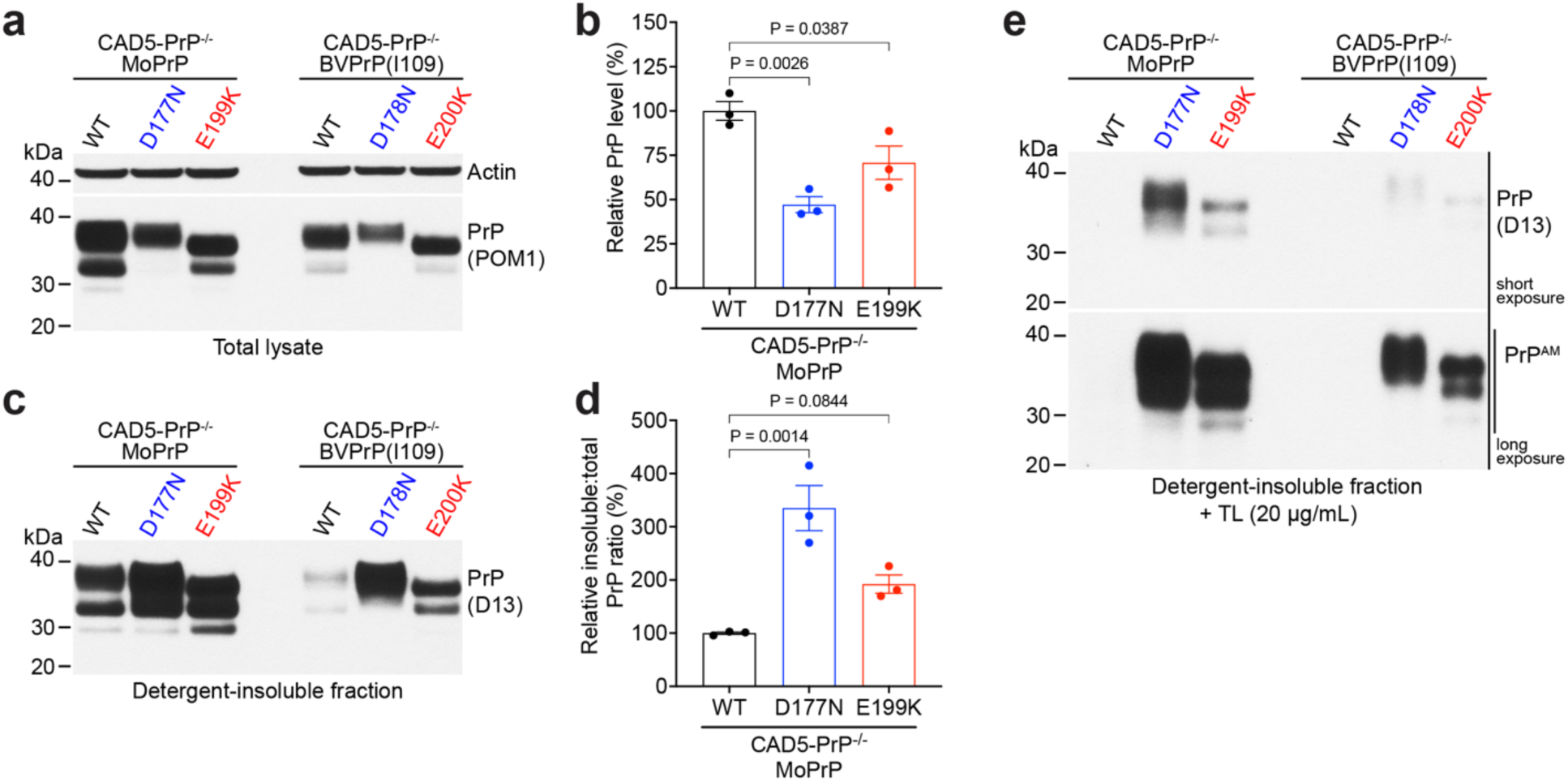
Stably transfected CAD5-PrP^-/-^ cells expressing D177N- or E199K-mutant mouse PrP generate PrP^AM^. **a**) Representative immunoblot of total PrP levels in lysates from polyclonal lines of stably transfected CAD5-PrP^-/-^ cells expressing the indicated MoPrP constructs. The blot was reprobed with an antibody against actin. **b**) Quantification of total PrP levels in lysates from stably transfected CAD5-PrP^-/-^ cells. **c**) Representative immunoblot of detergent-insoluble PrP levels in lysates from CAD5-PrP^-/-^ cells stably transfected with the indicated MoPrP constructs. **d**) Quantification of detergent-insoluble PrP levels (normalized to total PrP levels) in lysates from stably transfected CAD5-PrP^-/-^ cells. **e**) Representative immunoblot of detergent-insoluble, thermolysin-resistant PrP species (“PrP^AM^”) in lysates from polyclonal lines of stably transfected CAD5-PrP^-/-^ cells expressing the indicated MoPrP or BVPrP(I109) constructs. In panels b and d, data are mean ± SEM from n = 3 independent samples, and statistical significance was assessed using one-way ANOVA followed by Dunnett’s multiple comparisons test.

To understand how *PRNP* mutations affect the stability of BVPrP(I109), we generated monoclonal CAD-PrP^-/-^ cell lines expressing either WT, D178N-mutant, or E200K-mutant BVPrP(I109). This strategy allowed us to select clones with approximately matched levels of PrP expression for WT or mutant BVPrP(I109) and, as expected, total PrP levels were higher in the monoclonal lines compared to the polyclonal lines (**Fig. 3a**). As in the polyclonal lines, detergent-insoluble and TL-resistant PrP^AM^ species were selectively produced in the cells expressing either D178N- or E200K-mutant BVPrP(I109) (**Fig. 3b**). Even when using a lower concentration of TL (5 µg/mL), PrP^AM^ was only present in lysates from monoclonal cells expressing mutant BVPrP(I109). To assess the relative stability of WT vs. mutant BVPrP(I109) in the monoclonal lines, we treated cells with cycloheximide (CHX) to shut off protein synthesis and then measured PrP levels at different timepoints post-CHX treatment. D178N-mutant BVPrP(I109) turned over more rapidly than either WT BVPrP(I109) or E200K-mutant BVPrP(I109) (**Fig. 3c, d**). The half-lives of WT, D178N-mutant, and E200K-mutant BVPrP(I109) in CAD5-PrP^-/-^ cells were calculated to be ∼18, ∼5, and ∼15 hours, respectively. Thus, the D178N mutation strongly destabilizes BVPrP(I109).

**Figure 3.**
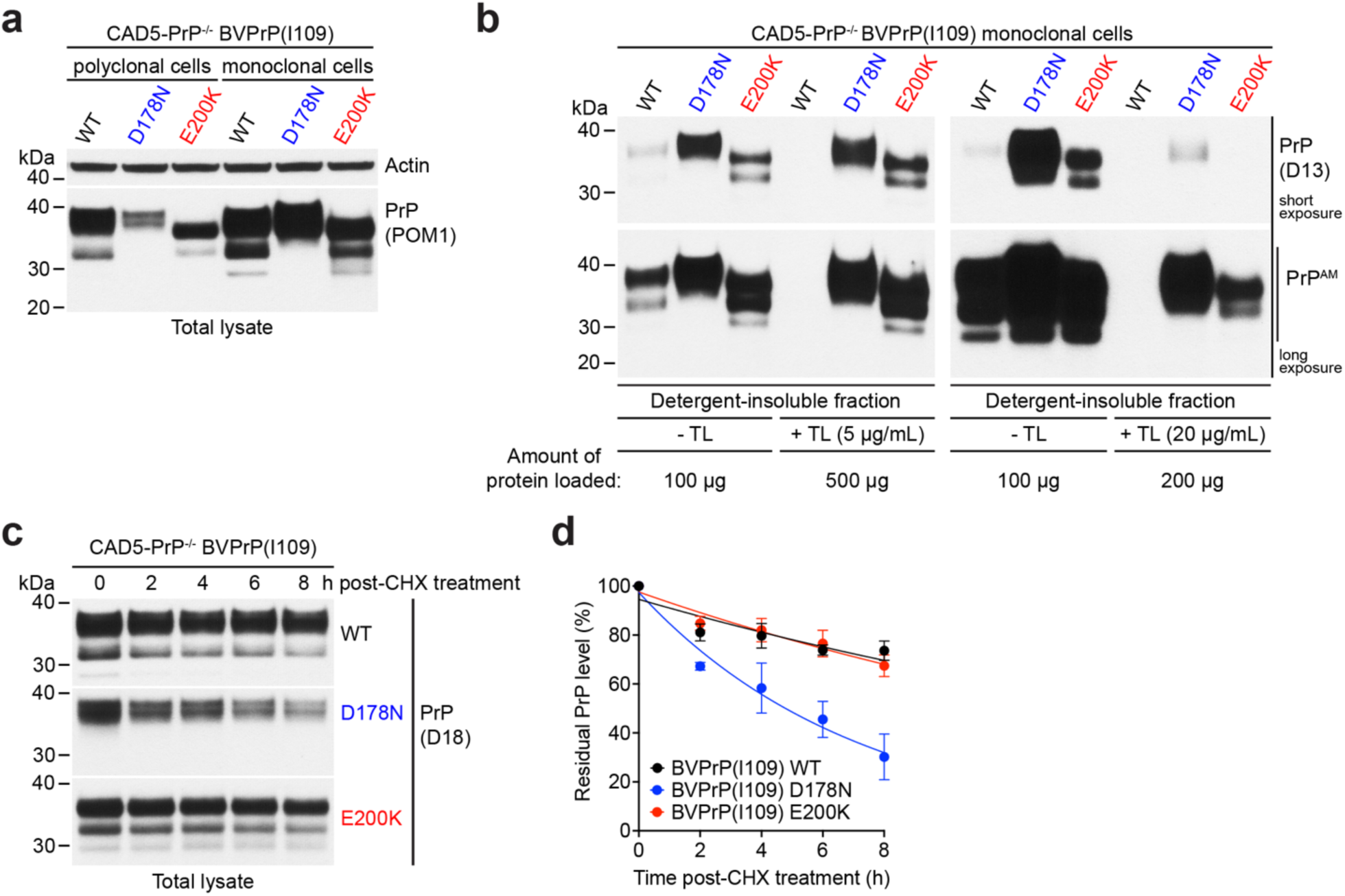
Destabilizing effect of the D178N mutation in monoclonal CAD5-PrP^-/-^ cell lines stably expressing bank vole PrP. **a**) Representative immunoblot of total PrP levels in lysates from polyclonal and monoclonal CAD5-PrP^-/-^ cells stably transfected with the indicated BVPrP(I109) constructs. The blot was reprobed with an antibody against actin. **b**) Representative immunoblots of detergent-insoluble PrP species in lysates from monoclonal stably transfected CAD5-PrP^-/-^ cells that were either treated (+ TL) or not treated (- TL) with thermolysin. The blot on the left depicts samples treated with 5 µg/mL TL whereas the blot on the right depicts samples treated with 20 µg/mL TL. **c**) Immunoblots of residual PrP levels in lysates from stably transfected monoclonal CAD5-PrP^-/-^ lines expressing either WT, D178-mutant, or E200K-mutant BVPrP(I109) treated with cycloheximide (CHX) for the indicated amounts of time. **d**) Quantification of residual PrP levels in the monoclonal cell lines following treatment with CHX for the indicated amounts of time. Data are mean ± SEM for n = 3 independent experiments.

Recently, we demonstrated that a subset of knock-in mice expressing physiological levels of D178N- or E200K-mutant BVPrP(I109), respectively termed kiBVI^D178N^ and kiBVI^E200K^, develop spontaneous prion disease with an onset between 400 and 600 days of age (68). The brains of spontaneously sick kiBVI^D178N^ and kiBVI^E200K^ mice selectively exhibit PrP species that are resistant to digestion with a high concentration of TL (100 µg/mL). Consistent with previous results (68), steady-state PrP levels were significantly lower in the brains of young (3-month-old) asymptomatic kiBVI^D178N^ mice and slightly lower in the brains of kiBVI^E200K^ mice compared to kiBVI^WT^ mice expressing WT BVPrP(I109) (**Fig. 4a, b**). Like what we observed in cells, relative levels of detergent-insoluble PrP species were markedly elevated in brains from young kiBVI^D178N^ and kiBVI^E200K^ compared to kiBVI^WT^ mice (**Fig. 4c, d**). To determine if PrP^AM^ is present in the brains of young, asymptomatic knock-in mice, brain homogenates were digested with 20 µg/mL TL. TL-resistant PrP species were observed in the brains of young kiBVI^D178N^ and kiBVI^E200K^ mice but not in the brains of kiBVI^WT^ mice (**Fig. 4e**). Hence, PrP^AM^ is present in the brains of knock-in mice expressing mutant BVPrP(I109) approximately one year prior to the typical onset of prion disease.

**Figure 4.**
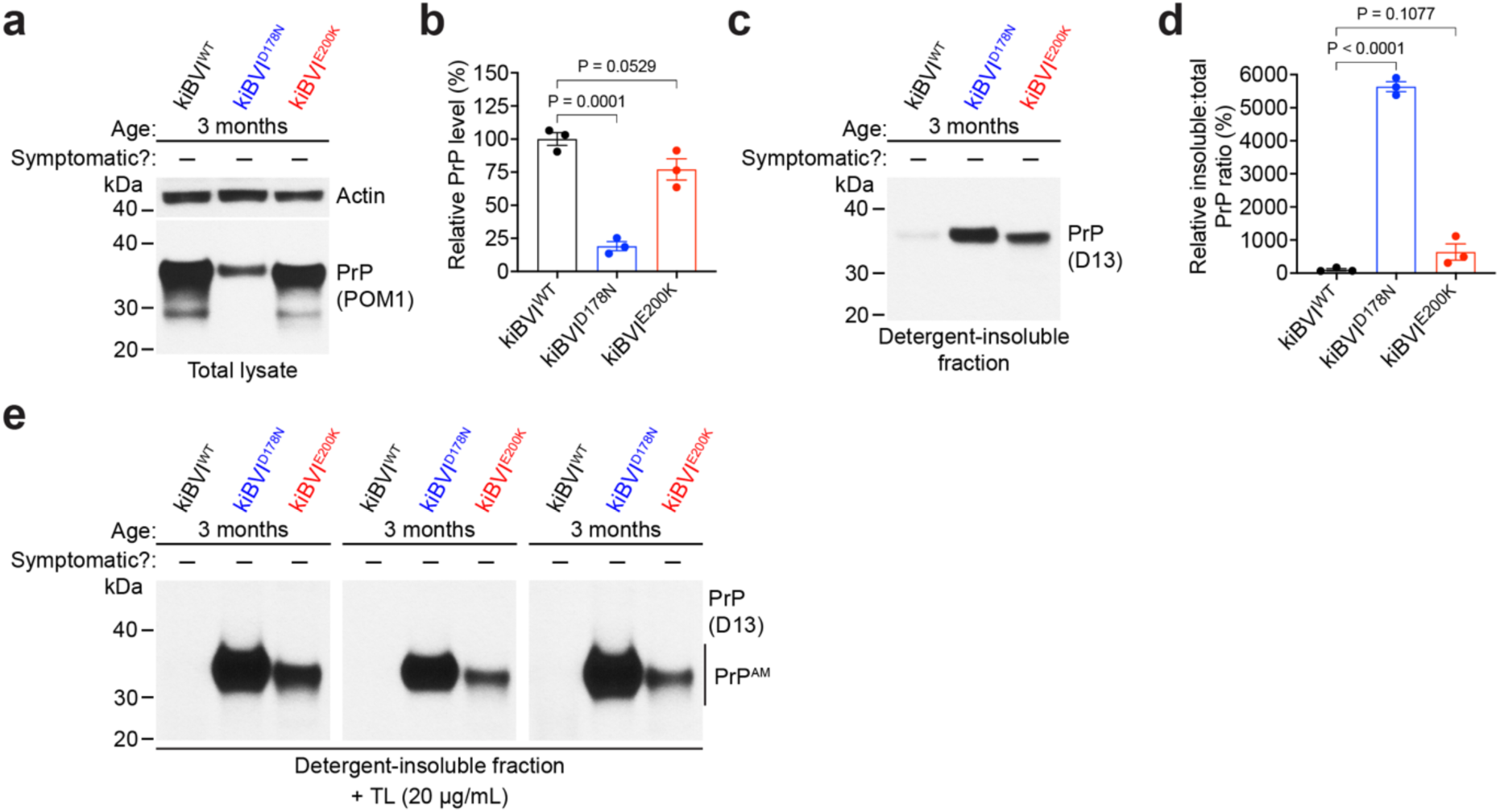
The brains of young, asymptomatic knock-in mice expressing mutant bank vole PrP contain PrP^AM^. **a**) Representative immunoblot of total PrP levels in brain homogenates from 3-month-old kiBVI^WT^, kiBVI^D178N^, and kiBVI^E200K^ mice. The blot was reprobed with an antibody against actin. **b**) Quantification of total PrP levels in brain homogenates from 3-month-old kiBVI^WT^, kiBVI^D178N^, and kiBVI^E200K^ mice. **c**) Representative immunoblot of detergent-insoluble PrP species in brain homogenates from 3-month-old kiBVI^WT^, kiBVI^D178N^, and kiBVI^E200K^ mice. **d**) Quantification of detergent-insoluble PrP levels (normalized to total PrP levels) in brain homogenates from 3-month-old kiBVI^WT^, kiBVI^D178N^, and kiBVI^E200K^ mice. **e**) Representative immunoblots of detergent-insoluble, thermolysin-resistant PrP species (“PrP^AM^”) in brain homogenates from 3 independent sets of 3-month-old kiBVI^WT^, kiBVI^D178N^, and kiBVI^E200K^ mice. In panels b and d, data are mean ± SEM from n = 3 independent samples, and statistical significance was assessed using one-way ANOVA followed by Dunnett’s multiple comparisons test.

### PrP^AM^ production is enhanced by the prion disease protective G127V mutation

We reasoned that PrP^AM^ could either represent a direct precursor of classical or atypical PrP^Sc^ species observed during clinical prion disease or an off-pathway misfolded species. To distinguish between the two possibilities, we first asked whether the G127V mutation might counteract the production of PrP^AM^. The G127V mutation, which was discovered in individuals seemingly resistant to kuru (81), interferes with prion replication and prevents or delays prion disease even when inserted into permissive substrates such as BVPrP (74, 82–84). If PrP^AM^ is a direct precursor of PrP^Sc^, the G127V mutation might be predicted to decrease or abolish PrP^AM^ formation. The G127V mutation was inserted into the sequence of WT, D178N-mutant, and E200K-mutant BVPrP(I109) (**Fig. 5a**), and then polyclonal lines of CAD5-PrP^-/-^ cells stably expressing G127V-containing BVPrP(I109) variants were generated. For WT and E200K-mutant BVPrP(I109) the G127V mutation had no effect on PrP expression levels (**Fig. 5b, c**). However, addition of the G127V mutation significantly decreased levels of D178N-mutant BVPrP(I109) expression in cells. This may potentially be explained by a decreased stability of D178N-mutant BVPrP(I109) containing the G127V mutation, as assessed following CHX treatment (**Fig. S5a, b**). Cell-surface PrP expression was detectable in CAD5-PrP^-/-^ cells stably expressing WT, D178N-mutant, or E200K-mutant BVPrP(I109), and cell surface localization was unaltered by the G127V mutation (**Fig. S5c**).

**Figure 5.**
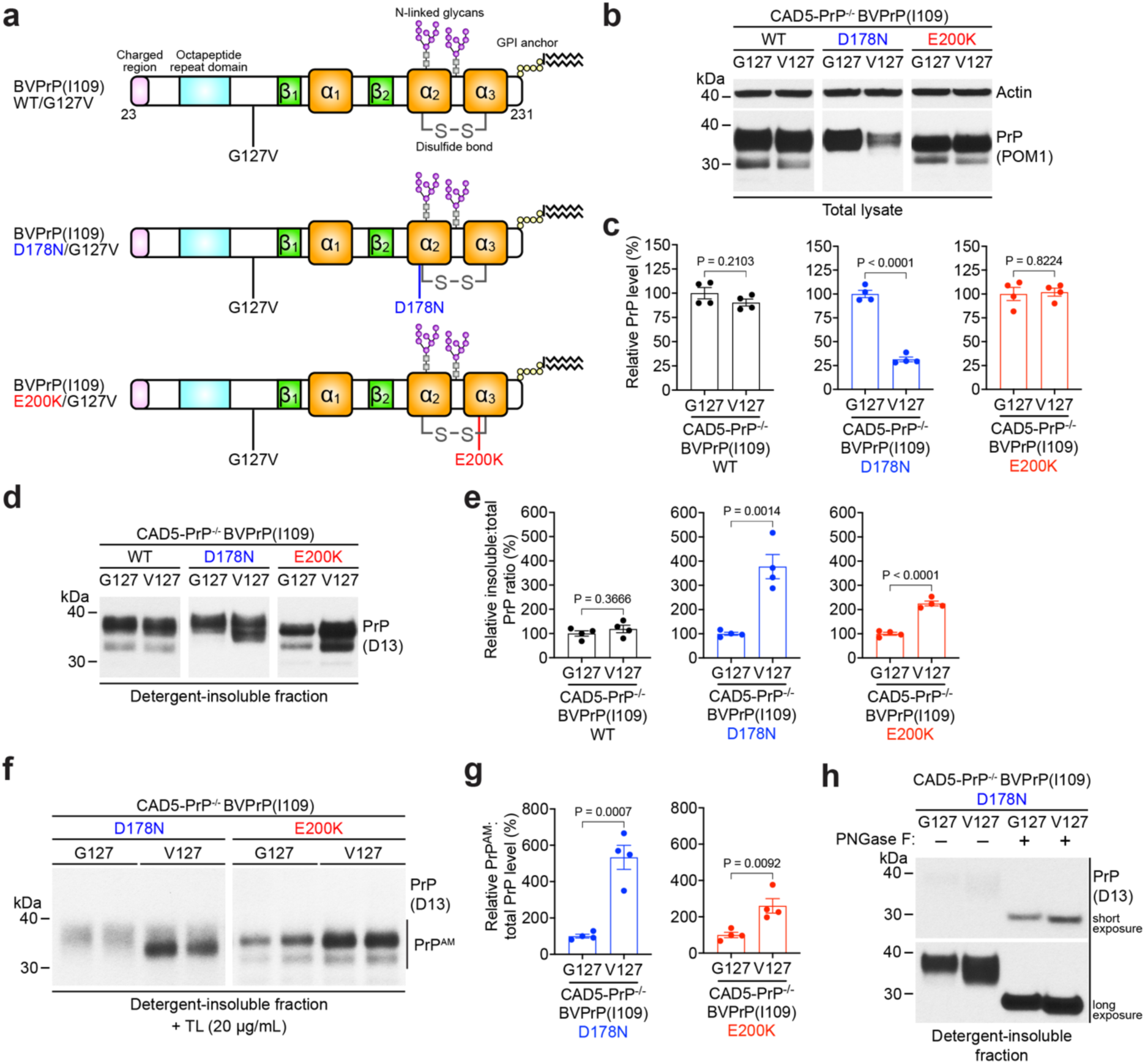
The prion disease protective G127V mutation enhances PrP^AM^ production in CAD5-PrP^-/-^ cells expressing mutant bank vole PrP. **a**) Schematic structures of WT, D178N, and E200K BVPrP(I109) containing the G127V mutation. **b**) Representative immunoblots of total PrP levels in lysates from CAD5-PrP^-/-^ cells stably transfected with the indicated BVPrP(I109) constructs containing either G127 or V127. The blots were reprobed with an antibody against actin. **c**) Quantification of total PrP levels in lysates from stably transfected CAD5-PrP^-/-^ cells. **d**) Representative immunoblots of detergent-insoluble PrP levels in lysates from CAD5-PrP^-/-^ cells stably transfected with the indicated constructs. **e**) Quantification of detergent-insoluble PrP levels (normalized to total PrP levels) in lysates from stably transfected CAD5-PrP^-/-^ cells. **f**) Representative immunoblots of detergent-insoluble, thermolysin-resistant PrP species (“PrP^AM^”) in lysates from stably transfected CAD5-PrP^-/-^ cells. **g**) Quantification of PrP^AM^ levels (normalized to total PrP levels) in lysates from stably transfected CAD5-PrP^-/-^ cells. **h**) Immunoblot of detergent-insoluble PrP species in lysates from cells expressing D178N-mutant BVPrP(I109) containing either G127 or V127. Following ultracentrifugation, samples were either treated with PNGase F or left untreated. In panels c, e, and g, data are mean ± SEM from n = 4 independent samples, and statistical significance was assessed using two-tailed unpaired t tests.

While the G127V mutation had no effect on the relative amount of detergent-insoluble PrP species in cells expressing WT BVPrP(I109), the addition of G127V significantly increased the relative amounts of detergent-insoluble PrP in cells expressing D178N- or E200K-mutant BVPrP(I109) (**Fig. 5d, e**). Levels of TL-resistant PrP^AM^ species were significantly enhanced by the G127V mutation in cells expressing either D178N- or E200K-mutant BVPrP(I109) (**Fig. 5f, g**). Addition of the G127V mutation resulted in an ∼5-fold increase in PrP^AM^ levels for D178N and an ∼2.5-fold increase for E200K. The G127V mutation also caused *de novo* production of potential PrP^AM^-like species in cells expressing WT BVPrP(I109), although the effect was less dramatic (**Fig. S6**). For D178N, the G127V-mediated increase in detergent-insoluble species and PrP^AM^ levels appeared to predominantly result from the presence of a PrP species with a lower apparent molecular weight (**Fig. 5d, f**). To check if this might be due to differential N-glycosylation, the detergent-insoluble fraction from cells expressing D178N-mutant BVPrP(I109) with or without the G127V mutation was treated with PNGase F. After enzymatic deglycosylation, the molecular weights of detergent-insoluble D178N-mutant BVPrP(I109) species containing either G127 or V127 were similar (**Fig. 5h**), suggesting that the G127V mutation preferentially increases production of misfolded monoglycosylated D178N-mutant BVPrP(I109) species. Thus, contrary to our initial prediction, the G127V mutation appears to potentiate the misfolding-promoting effects of the D178N and E200K mutations in BVPrP(I109).

### PrP^AM^ is unaffected by treatment with anti-prion small molecules

Next, we assessed whether small molecule anti-prion compounds have any effect on PrP^AM^ levels in cultured cells expressing mutant BVPrP(I109). Both IND24 and Anle138b decrease levels of PK-resistant PrP in prion-infected cultured cells, and treatment of prion-infected mice with IND24 or Anle138b delays the onset of prion disease (74, 85–87). If PrP^AM^ is a precursor of PrP^Sc^, treatment with anti-prion small molecules may dissuade PrP^AM^ formation. First, we confirmed that treatment of CAD5-PrP^-/-^ BVPrP(I109) cells infected with BVPrP-adapted RML prions for 3 days with either IND24, Anle138b, or both compounds simultaneously results in a significant decrease in PK-resistant PrP levels (**Fig. 6a, b**). To assess whether the compounds have any effect on PrP^AM^ levels, CA5-PrP^-/-^ cells stably expressing either WT, D178N-mutant, or E200K-mutant BVPrP(I109) were treated with IND24, Anle13b, or both compounds for 3 days. Neither compound, either alone or in combination, had any effect on total PrP levels (**Fig. 6c, d**) or the relative amounts of detergent-insoluble PrP species in cells (**Fig. 6e, f**). Similarly, levels of PrP^AM^ were unaffected by treatment with the anti-prion compounds (**Fig. 6g, h**).

**Figure 6.**
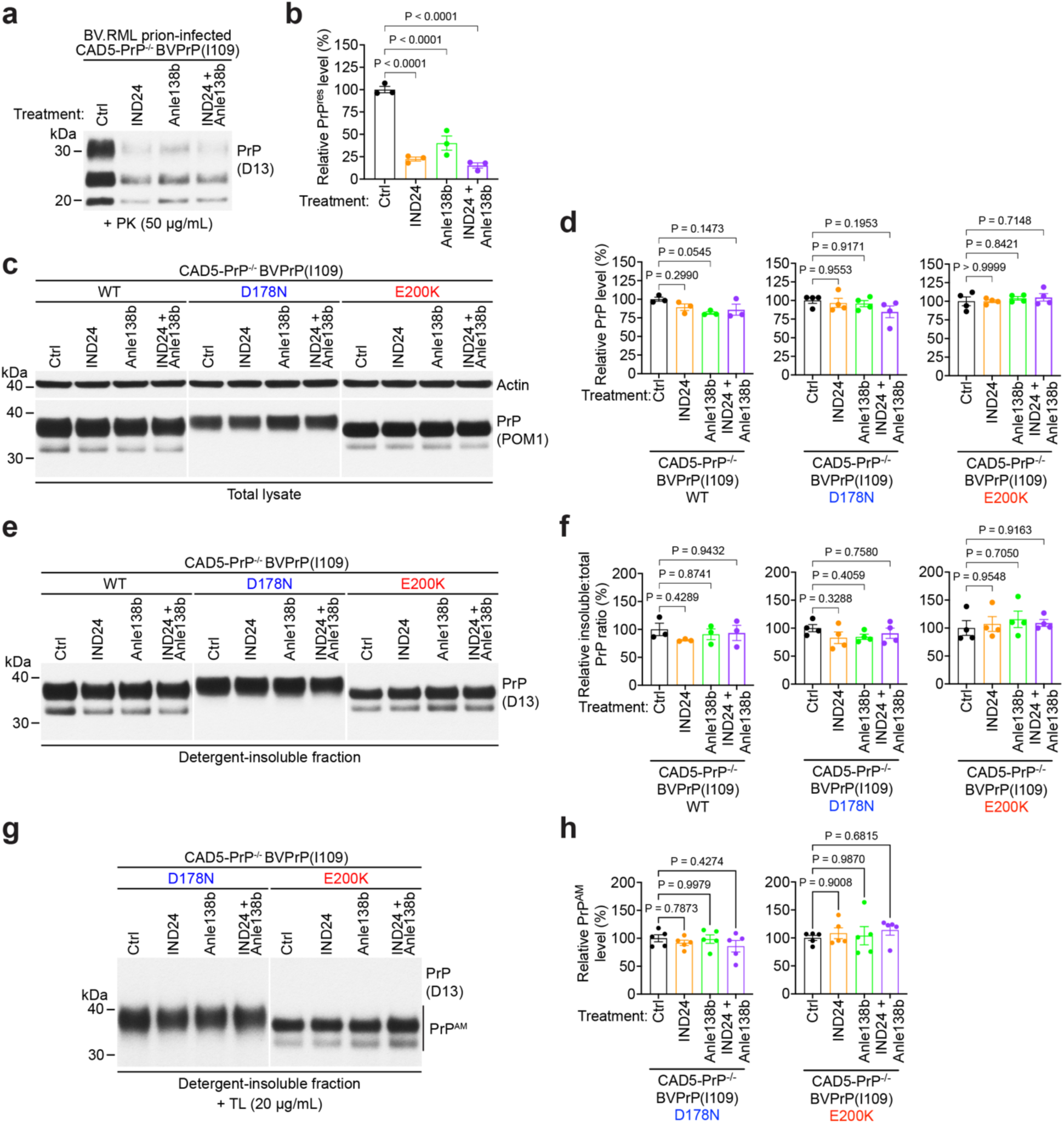
The anti-prion small molecules IND24 and Anle138b have no effect on PrP^AM^ levels in CAD5-PrP^-/-^ cells expressing mutant bank vole PrP. **a**) Representative immunoblot of PK-resistant PrP levels in lysates from monoclonal CAD5-PrP^-/-^ BVPrP(I109) cells infected with BV.RML prions and treated with the indicated anti-prion compounds for 3 days. **b**) Quantification of PK-resistant PrP levels in cells treated with anti-prion compounds (n = 3 independent replicates per treatment). **c**) Representative immunoblots of total PrP levels in lysates from CAD5-PrP^-/-^ cells stably transfected with the indicated BVPrP(I109) constructs and treated with the indicated anti-prion compounds for 3 days. The blots were reprobed with an antibody against actin. **d**) Quantification of total PrP levels in lysates from cells treated with anti-prion compounds (n = 3 independent replicates for WT, n = 4 independent replicates for D178N and E200K). **e**) Representative immunoblots of detergent-insoluble PrP species in lysates from CAD5-PrP^-/-^ cells stably transfected with the indicated BVPrP(I109) constructs and treated with the indicated anti-prion compounds for 3 days. **f**) Quantification of detergent-insoluble PrP levels in lysates from cells treated with anti-prion compounds (n = 3 independent replicates for WT, n = 4 independent replicates for D178N and E200K). **g**) Representative immunoblots of detergent-insoluble, thermolysin-resistant PrP species (“PrP^AM^”) in lysates from stably transfected CAD5-PrP^-/-^ cells expressing D178N- or E200K-mutant BVPrP(I109) and treated with anti-prion compounds for 3 days. **h**) Quantification of PrP^AM^ levels in lysates from cells treated with anti-prion compounds (n = 5 independent replicates). In panels b, d, f, and h, data are mean ± SEM, and statistical significance was assessed using one-way ANOVA followed by Dunnett’s multiple comparisons test.

### Cells harboring PrP^AM^ are largely resistant to prion infection

If PrP^AM^ is on-pathway for prion formation, then the presence of PrP^AM^ in cells should facilitate prion infection. First, we challenged the polyclonal lines of CAD5-PrP^-/-^ cells expressing either WT, D178N-mutant, or E200K-mutant BVPrP(I109) with four strains of BVPrP-adapted prions: BV.RML, BV.22L, BV.263K, and BV.HY (63). BV.RML and BV.22L are mouse prion strains that have been passaged in cells expressing BVPrP(I109). Likewise, BV.263K and BV.HY are hamster prion strains that have been passaged in cells expressing BVPrP(I109). Cells were exposed to prion-containing cell homogenate for 3 days and then passaged 5 times to remove any residual prion inoculum. While cells expressing WT BVPrP(I109) could become readily infected with all four strains of BVPrP-adapted prions, as judged by the presence of PK-resistant PrP in cell lysates, cells expressing D178N-mutant BVPrP(I109) were resistant to prion infection (**Fig. 7a**). Small amounts of PK-resistant PrP were observed following infection of cells expressing E200K-mutant BVPrP following challenge with BV.RML or BV.22L prions, but not with BV.263K or BV.HY prions.

**Figure 7.**
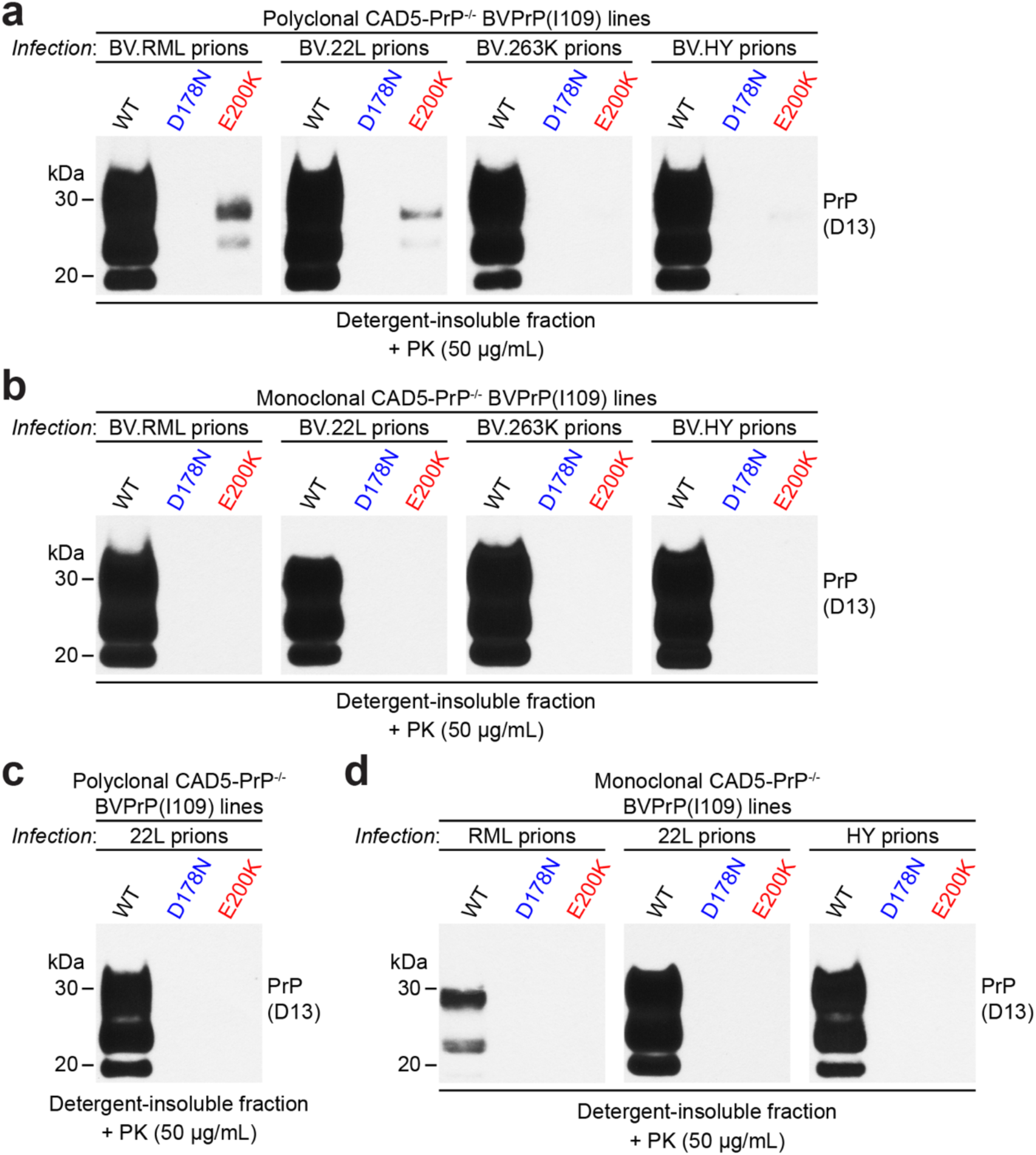
The D178N and E200K mutations restrict prion infection. Representative immunoblots of detergent-insoluble PK-resistant PrP levels in lysates from polyclonal (a, c) or monoclonal (b, d) lines of stably transfected CAD5-PrP^-/-^ cells expressing either WT, D178N-mutant, or E200K-mutant BVPrP(I109) at passage 5 following infection with the indicated BVPrP-adapted prion strains (a, b) or with mouse 22L, mouse RML, or hamster HY prions (c, d).

PrP^C^ levels are lower in the polyclonal CAD5-PrP^-/-^ lines expressing D178N- or E200K-mutant BVPrP(I109) compared to lines expressing WT BVPrP(I109) (**Fig. 1a**). Thus, the reduced ability to infect the polyclonal mutant BVPrP-expressing cells could be due to lower PrP^C^ levels rather than a direct effect of the mutations on prion replication. To address this issue, the monoclonal cell lines expressing WT or mutant BVPrP(I109), which have roughly matched PrP^C^ expression levels (**Fig. 3a**), were challenged with prions. Whereas the WT monoclonal line was similarly susceptible to the same four BVPrP-adapted prion strains, the monoclonal D178N and E200K lines were almost completely resistant to prion infection (**Fig. 7b, S7**). We also checked whether the polyclonal and monoclonal BVPrP(I109) cell lines are susceptible to the original, non-BVPrP-adapted prion strains. Polyclonal lines expressing WT BVPrP(I109), but not D178N-mutant or E200K-mutant BVPrP(I109), were directly susceptible to infection with mouse 22L prions (**Fig. 7c**). Likewise, monoclonal lines expressing WT BVPrP(I109), but not D178N- or E200K-mutant BVPrP(I109), were directly susceptible to infection with mouse RML and 22L prions as well as hamster HY prions (**Fig. 7d**). Thus, instead of promoting prion infection, the D178N and E200K mutations prevent or hinder prion infection in cultured cells.

### Cells harboring PrP^AM^ do not exhibit prion seeding activity

Lastly, we assessed whether cell homogenates containing PrP^AM^ exhibit prion seeding activity, as may be expected if PrP^AM^ is a precursor of PrP^Sc^. We rationalized that if PrP^AM^ can act as a seed and propagate, then exposure of cells to PrP^AM^-containing homogenate ought to increase the relative amount of PrP^AM^ in the recipient cells. Polyclonal lines of stably-transfected cells expressing either WT or D178N-mutant BVPrP(I109) were exposed to homogenate from CAD5-PrP^-/-^ BVPrP(I109) cells containing either no PrP^AM^ (WT), moderate levels of PrP^AM^ (D178N), or high levels of PrP^AM^ (D178N/G127V) and then passaged 3 times (**Fig. 8a**). As expected, total PrP levels in both the WT and D178N lines remained unaltered regardless of the treatment (**Fig. 8b**). There was no obvious increase in either detergent-insoluble PrP levels (**Fig. 8c**) or PrP^AM^ levels (**Fig. 8d**) in D178N cells exposed to D178N or D178N/G127V homogenate. Likewise, treatment with PrP^AM^ -containing samples did not induce *de novo* formation of PrP^AM^ in cells expressing WT BVPrP(I109) (**Fig. 8d**). We also checked whether PrP^AM^-containing cell homogenates exhibit prion seeding activity in the real-time quaking-induced conversion (RT-QuIC) assay. This assay measures the ability of PrP^Sc^ to stimulate the aggregation of recombinant PrP, which is assessed by monitoring Thioflavin T (ThT) fluorescence over time (88). We used recombinant bank vole PrP (M109 variant) as the substrate for RT-QuIC, as it has been shown to function as permissive substrate for many different classical and atypical prion strains, including the prions generated spontaneously in the brains of kiBVI^D178N^ and kiBVI^E200K^ mice (62, 68). As a positive control we used homogenate from CAD5-PrP^-/-^ BVPrP(I109) cells infected with BV.RML prions and as a negative control we utilized homogenate from CAD5-PrP^-/-^ cells. While 4/5 replicates from the BV.RML prion infected cells were positive in the RT-QuIC assay, as judged by a reduced lag phase, there was no significant difference between the lag phases for samples from cells expressing D178N- or D178N/G127V-mutant BVPrP(I109) and samples from CAD-PrP^-/-^ cells (**Fig. 8e, f**). Therefore, samples containing PrP^AM^ do not exhibit prion seeding activity, either *in vitro* or in cells.

**Figure 8.**
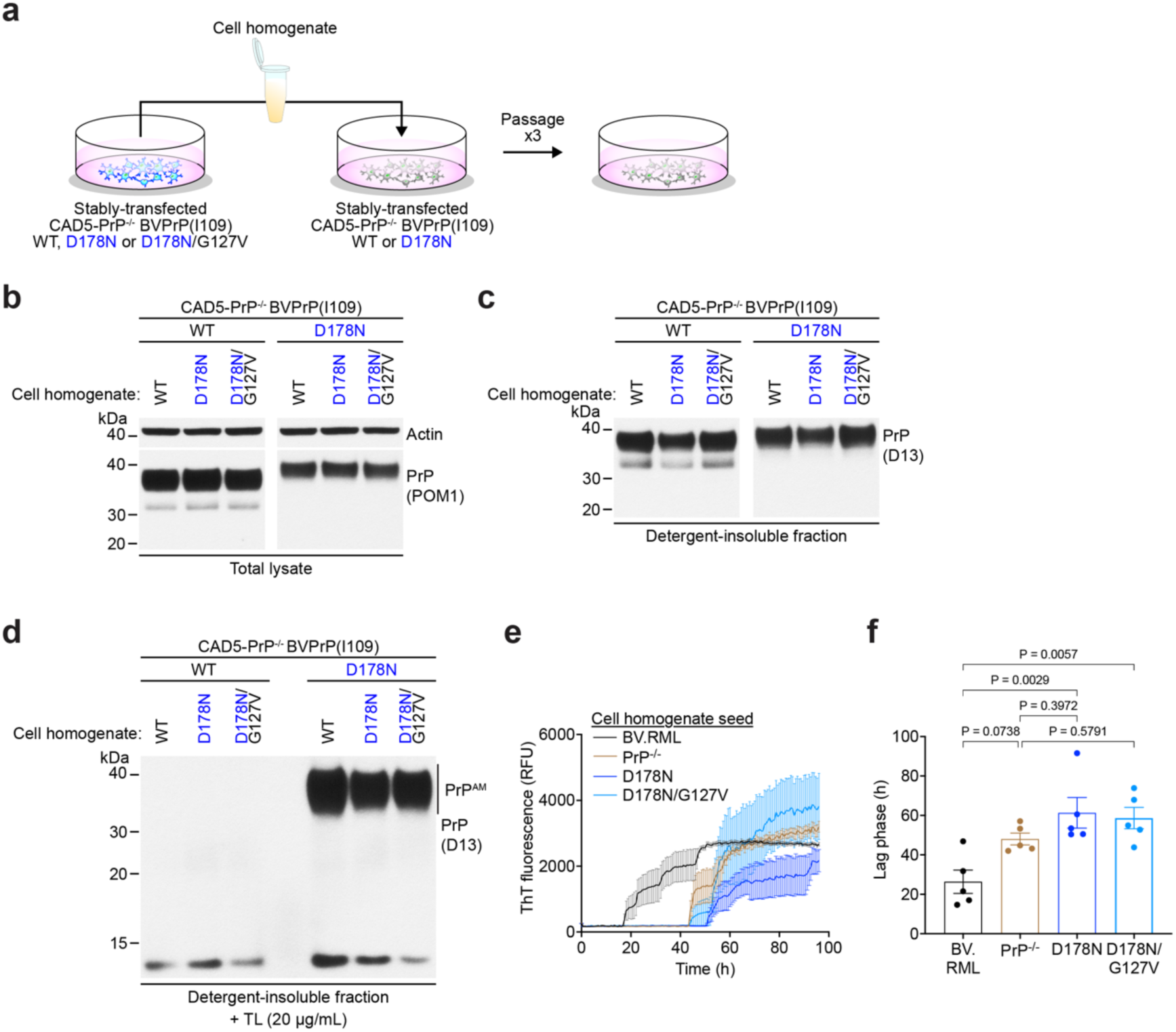
PrP^AM^-containing homogenates from CAD5-PrP^-/-^ cells expressing mutant bank vole PrP do not exhibit prion seeding activity. **a**) Schematic of experiment in which homogenates from stably transfected CAD5-PrP^-/-^ cells expressing either WT, D178N-mutant, or D178N/G127V-mutant BVPrP(I109) were applied to cells expressing either WT or D178N-mutant BVPrP(I109) to check for PrP^AM^-induced seeding activity. **b**) Representative immunoblots of total PrP levels in lysates from CAD5-PrP^-/-^ cells expressing either WT or D178N-mutant BVPrP(I109) after 3 passages following treatment with homogenate from the indicated cell lines. The blots were reprobed with an antibody against actin. **c**) Representative immunoblots of detergent-insoluble PrP species in lysates from CAD5-PrP^-/-^ cells expressing either WT or D178N-mutant BVPrP(I109) after 3 passages following treatment with homogenate from the indicated cell lines. **d**) Representative immunoblots of detergent-insoluble, thermolysin-resistant PrP species (“PrP^AM^”) in lysates from CAD5-PrP^-/-^ cells expressing either WT or D178N-mutant BVPrP(I109) after 3 passages following treatment with homogenate from the indicated cell lines. **e**) Thioflavin T (ThT) fluorescence curves for RT-QuIC experiments using homogenates from cells expressing either D178N- or D178N/G127V-mutant BVPrP(I109) as the seed. Homogenate from CAD5-PrP^-/-^ cells was used as a negative control and homogenate from CAD5-PrP^-/-^ BVPrP(I109) cells infected with RML prions (BV.RML) was used as a positive control. Data are mean ± SEM from n = 5 independent replicates. **f**) Quantification of lag phases for the RT-QuIC experiments. Data are mean ± SEM from n = 5 independent replicates, and statistical significance was assessed using one-way ANOVA followed by Dunnett’s multiple comparisons test.

## Discussion

Here, we show that, in both cultured cells and mice, the D178N and E200K *PRNP* mutations lead to formation of an alternatively misfolded PrP species, which we term PrP^AM^, that is TL-resistant but does not exhibit any detectable prion seeding activity. Since PrP^AM^ is both detergent-insoluble and TL-resistant, it seems likely that PrP^AM^ is comprised of PrP aggregates, but the precise assembly state of PrP^AM^ is currently unknown. TL cleavage occurs prior to bulky hydrophobic amino acids, which are notably absent within the disordered N-terminal domain of PrP (68), implying that hydrophobic residues within the C-terminal domain undergo conformational re-arrangement and become less accessible in PrP^AM^. As PrP^AM^ is resistant to modest concentrations of TL but sensitive to PK digestion, this could suggest that it represents a smaller, potentially oligomeric PrP assembly. Although smaller PrP assemblies have been reported to be more toxic than their larger counterparts (89, 90), PrP^AM^ itself is unlikely to be highly toxic since it is present in the brains of young healthy kiBVI^D178N^ and kiBVI^E200K^ mice. Instead, we hypothesize that PrP^AM^ is an innocuous misfolded species that may exert protective properties towards preventing mutant PrP from becoming PrP^Sc^. The existence of a protective PrP assembly that actively dissuades PrP^Sc^ formation provides a potential explanation for why genetic prion disorders typically manifest later in life despite the production of mutant PrP from birth.

Although both the D178N and E200K mutations generate PrP^AM^, the protective effects of this species must eventually wane since individuals with these *PRNP* mutations ultimately develop prion disease with nearly complete penetrance (91). Potential relationships between PrP^AM^ and PrP^Sc^ will require further investigation. It is conceivable that the formation of both PrP^AM^ and PrP^Sc^ could be enhanced by slowed folding kinetics that alter the exposure of aggregation-prone hydrophobic residues within the protein. The rate-limiting step in the sporadic and genetic prion diseases is believed to be the spontaneous formation of a stable PrP^Sc^ seed that can self-propagate (92), although the precise mechanistic basis for prion formation remains to be elucidated. It is possible that PrP^AM^ may exert its protective effects by actively blocking one or more steps during PrP^Sc^ seed generation, a process that may become less efficient with ageing due to an increase in seed production stemming from a reduction in brain proteostatic machinery (93). Alternatively, PrP^AM^ may interfere with prion propagation by preventing association of PrP^Sc^ with PrP^C^ and thus preventing elongation of PrP^Sc^ aggregates.

A fundamental question is whether PrP^AM^ itself is on-pathway or off-pathway for prion formation. Because PrP^AM^ appears to lack prion seeding activity, is promoted by the prion protective G127V mutation, and is present in mutant PrP-expressing cells that are resistant to prion infection, our findings are more consistent with PrP^AM^ being off-pathway. However, based on studies in knock-in mice that develop spontaneous prion disease, it has been proposed that TL-resistant PrP aggregates may form first and then conformationally evolve into PK-resistant aggregates (68). Thus, it is possible that PrP^AM^, which is resistant to lower concentrations of TL, undergoes a structural transition to form aggregates that are resistant to higher concentrations of TL, and then eventually yields PK-resistant PrP^Sc^ later during the disease course. Therefore, if PrP^AM^ is on-pathway for prion formation, it would most likely represent a precursor that only rarely converts to a disease-associated form. Cofactor molecules are required for WT, but not D178N-mutant or E200K-mutant PrP substrates to form PrP^Sc^ *in vitro* (94). *In vivo*, kiBVI^E200K^ mice form PrP^Sc^ more readily when challenged with cofactor-less recombinant PrP fibrils than mice expressing WT BVPrP(I109) (95). These results raise the possibility that PrP^AM^ could be both an early protective species as well as a precursor to PrP^Sc^.

Cells expressing D178N- or E200K-mutant BVPrP(I109) were largely resistant to infection with several prion strains, which supports our hypothesis that PrP^AM^ dissuades prion replication. Consistent with our findings in cells, knock-in and transgenic mice expressing D177N-mutant MoPrP take longer to develop prion disease or do not develop disease at all following prion infection (42, 96). To the best of our knowledge, prion challenge of mice expressing E200K-mutant PrP has not yet been conducted. If the D178N- and E200K mutations simply increase PrP^Sc^ formation by either destabilizing PrP^C^ or stabilizing PrP^Sc^, one would have expected that the mutations would promote prion replication in cells. Instead, we hypothesize that it is the mutation-induced production of PrP^AM^ that hinders prion replication. Indeed, PrP^AM^ was more efficiently produced in cells expressing D178N-mutant BVPrP(I109) than in cells expressing E200K-mutant BVPrP(I109), and the D178N cells appeared to be completely resistant to prion infection whereas prion infection was only impeded in the E200K cells. However, an alternate explanation that must be considered is that the D178N and E200K mutations introduce a transmission barrier that prevents infection with the prion strains we tested, even when introduced into a permissive substrate such as BVPrP. In prion disease, transmission barriers are dictated by molecular compatibility between the PrP^Sc^ seed and the PrP^C^ substrate and are strongly influenced by differences in amino acid sequences between PrP^Sc^ and PrP^C^ (97). When D178N-mutant PrP is used as a substrate for protein misfolding cyclic amplification, a cell-free technique for studying prion replication, efficient production of PrP^Sc^ occurs following seeding with RML prions (46, 96). This implies that the inhibitory effect of the D178N mutation on RML prion replication requires intact cells, which favors the notion that it is mediated by production of PrP^AM^ rather than a transmission barrier. Nonetheless, further studies will be required to clarify this issue.

The mechanistic basis for how the G127V mutation impedes prion replication remains unknown. *In vitro*, the mutation interferes with the ability of recombinant PrP to polymerize into fibrils (74, 98, 99), although the relevance of recombinant PrP fibrils to prion replication *in vivo* remains unclear. Others have suggested that the G127V mutation may reduce the cytotoxicity of PrP aggregates or hinder PrP dimerization (100–102), although the latter effect was not observed in a cellular environment (103). Our findings suggest an alternative possibility for the protective nature of G127V, namely that it promotes formation of PrP^AM^ and thus shifts prion assembly towards a pathway that does not directly lead to PrP^Sc^ production. As G127V-mutant PrP can also prevent the formation of prions from WT PrP when the two molecules are co-expressed (82–84), this may suggest that PrP^AM^ derived from mutant PrP can help to prevent WT PrP from misfolding into PrP^Sc^. This provides a potential explanation for why in most cases of genetic prion disease, despite both WT and mutant PrP being produced simultaneously, only mutant PrP tends to be present in PrP^Sc^ (104, 105).

Consistent with the notion that PrP^AM^ is distinct from PrP^Sc^, PrP^AM^ levels were unaffected by treatment of mutant PrP-expressing cells with small molecules known to reduce PrP^Sc^ levels. The kinetics of spontaneous disease onset in kiBVI^D178N^ and kiBVI^E200K^ mice, which produce PrP^AM^ in their brains, is unaltered by treatment with the same anti-prion small molecules (IND24 and Anle138b) (106). Furthermore, Anle138b treatment does not increase survival in knock-in mice expressing either D177N- or E199K-mutant MoPrP, which would be predicted to harbor PrP^AM^ based on our findings in cultured cells (107). This provides further evidence that PrP^AM^ is not a direct precursor of neurotoxic and infectious PrP assemblies.

There are several limitations to this study. While we know that both the D178N and E200K mutations generate PrP^AM^, we don’t know if PrP^AM^ formation is a general feature of all prion disease-causing *PRNP* mutations, especially those that cause GSS. Furthermore, while we have shown that PrP^AM^ can form when *PRNP* mutations are introduced into either bank vole or mouse PrP, it will need to be investigated whether PrP^AM^ is produced when a human PrP backbone is used. Finally, additional structural information, including a thorough analysis of the aggregation state of PrP^AM^ as well as its biophysical and biochemical properties, will likely be required to clarify the role of PrP^AM^ in prion disease. Nonetheless, if PrP^AM^ is indeed protective against prion replication and/or formation, identifying cellular factors that regulate PrP^AM^ production and clearance could reveal new therapeutic targets for treating prion disease.

## Experimental Procedures

### Materials and chemicals

The following antibodies were used in this study: anti-PrP antibodies POM1 (Sigma #MABN2285) and POM2 (Sigma #MABN2298) as well as the recombinant humanized anti-PrP Fabs HuM-D18 and HuM-D13 (respectively referred to as D18 and D13 for simplicity), and an anti-actin antibody (Sigma #A5060). D18 was produced in-house whereas D13 was a generous gift from Stanley Prusiner. Horseradish peroxidase-conjugated secondary antibodies were purchased from Bio-Rad (#172-1011, #172-1019) and ThermoFisher (#31414). IND24 and Anle138b were custom synthesized by Sundia (Sacramento, CA) and were dissolved in DMSO.

### Plasmid constructs

Expression plasmids encoding wild-type bank vole PrP (I109 variant) or mouse PrP were generated by cloning the respective *Prnp* open reading frames between the *BamH*I and *Xba*l sites of the vector pcDNA3. Plasmids encoding D178N- or E200K-mutant BVPrP(I109) as well as D177N- or E199K-mutant MoPrP were generated by site-direct mutagenesis. The G127V substitution was introduced into the bank vole PrP plasmids by performing site-directed mutagenesis. All constructs were verified by DNA sequencing.

### Cell culture

Murine CAD5 cells, which were obtained from Charles Weissmann, are a subclone of the catecholaminergic CAD line (69, 70). CAD5-PrP^-/-^ cells were generated by CRISPR/Cas9 gene editing of CAD5 cells (72), and all experiments utilized clone D6. Cells were cultured in growth medium [Opti-MEM medium (ThermoFisher #31985088) containing 10% (v/v) fetal bovine serum (ThermoFisher #12483020) and 2 mM GlutaMAX (ThermoFisher #35050061)]. To passage the cells, they were first washed with DPBS (ThermoFisher #14190144) and then treated with enzyme-free cell dissociation reagent (MilliporeSigma #S-014-B). The cells were incubated for 1-3 minutes at 37 °C, dissociated, and then plated in fresh growth medium. Murine Neuro2a (N2a) neuroblastoma cells (ATCC# CCL-131) were cultured in DMEM medium (ThermoFisher #11965118) containing 10% (v/v) FBS, and 2 mM GlutaMAX. N2a-PrP^-/-^ cells were generated by CRISPR/Cas9 gene editing of N2a cells (108), and all experiments utilized clone 9.21. N2a and N2a-PrP^-/-^ cells were passaged by washing with DPBS and then treatment with 0.05% Trypsin-EDTA (ThermoFisher # 25300054). All cell lines were cultured at 37°C in a 5% CO_2_ atmosphere at constant humidity. Cells seeded from cryovials were cultured under passage-matched conditions and compared across samples, and cells with a passage number below 10 were used for analysis.

### Generation of stably transfected cell lines

For generation of stably transfected CAD5-PrP^-/-^ lines, CAD5-PrP^-/-^ cells were seeded at a density of 4.0-5.0 x 10^5^ cells/well in a 6-well plate. The following day, 2.0-2.5 µg of plasmid DNA was mixed with 4.0-5.0 µL of Lipofectamine-2000 (ThermoFisher #11668019) in 200 µL of serum-free Opti-MEM medium, and the mixture was added to the cells and incubated for 24 h. For generation of stably transfected N2a-PrP^-/-^ lines, N2a-PrP^-/-^ cells were seeded at a density of 4.0 x 10^5^ cells/well in a 6-well plate. Two days later, 2.0 µg of plasmid DNA was mixed with 4.0 µL of Lipofectamine-2000 in 200 µL of serum-free Opti-MEM medium, and the mixture was added to the cells and incubated for 24 h. After 24 h post-transfection, the cells were washed with DPBS and dissociated using the enzyme-free dissociation reagent. The transfected cells were then transferred into a 10-cm dish and cultured in medium containing 1 mg/mL G418. The cells were selected for 14 days and then expanded. The G418-containing medium was changed every three days. Polyclonal lines of stably transfected CAD5-PrP^-/-^ cells were maintained in growth medium containing 0.2 mg/mL G418, which ensures maintenance of PrP^C^ expression but does not interfere with *de novo* prion infection (109). For generation of monoclonal cell lines, stably transfected cells were diluted to 0.5 cells per 100 μL medium and then plated in 96-well plates. The plates were monitored over 3 weeks for the presence of single-cell-derived clones. Individual colonies were amplified sequentially in 24-well, 12-well, and 6-well plates, and then expanded further. Cells were lysed and checked for PrP^C^ expression by immunoblotting. Clones with approximately matched levels of PrP^C^ expression for WT or mutant BVPrP(I109) were picked. Monoclonal cell lines were cultured in the growth medium that did not contain G418.

### Cell lysis and immunoblotting

Cultured cells were washed with ice-cold DPBS and then lysed using lysis buffer, which consists of 50 mM Tris-HCl pH 7.4, 150 mM NaCl, 0.5% (w/v) sodium deoxycholate, and 0.5% (v/v) NP-40. The cell lysate was incubated on ice for 30 min, and then centrifuged at 5,000x *g* for 10 min to remove the insoluble cell debris. Protein concentrations in the supernatant were quantified using bicinchoninic acid (BCA) assay (ThermoFisher #23227), and then samples were diluted into 1X Bolt LDS sample buffer (ThermoFisher #B0007) to normalize protein concentration across samples. Unless otherwise indicated, all immunoblot samples were prepared under non-reduced conditions in the absence of β-mercaptoethanol and were not boiled. For reduced conditions, 2.5% (v/v) β-mercaptoethanol was added and samples were boiled at 95 °C for 10 min. Samples were electrophoresed on Bolt Bis-Tris gels (ThermoFisher #NW00100BOX and #NW00102BOX) using the MES buffer system. Following SDS-PAGE, proteins were transferred to Immobilon-P PVDF membranes (Millipore #IPVH00010), which were then blocked with blocking buffer [5 % (w/v) skim milk in Tris-buffered saline containing 0.05 % (v/v) Tween-20 (TBST)]. Membranes were then incubated overnight at 4 °C with antibodies diluted in blocking buffer. The following primary antibodies were used: POM1 (1:3,000-5,000 dilution), POM2 (1:5,000 dilution), D18 (1:5,000 dilution), D13 (1:10,000 dilution), or actin (1:5,000-10,000 dilution). The following day, membranes were washed 3 times with TBST (5 min per wash) and then incubated with horseradish peroxidase-conjugated secondary antibodies diluted in blocking buffer for 1 h at room temperature. The membranes were again washed 6 times with TBST (10 min per wash). The blots were developed using Western Lightning ECL pro (Revvity #NEL122001EA) or SuperSignal™ West Dura Extended Duration Substrate (ThermoFisher #34075) and then chemiluminescent signal was captured by exposure to HyBlot CL x-ray film (Thomas Scientific #1141J52). Immunoblotting data were quantified using NIH ImageJ software.

### Mouse brain homogenization and detergent extraction

Homozygous kiBVI^WT^, kiBVI^D178N^, and kiBVI^E200K^ mice were maintained as previously described in accordance with guidelines set by the Canadian Council on Animal Care under protocols (AUP 4263.24 and 6322.6) approved by the University Health Network Animal Care Committee (68). Brains from healthy 3-month-old knock-in mice were collected and divided parasagitally; the left-hemisphere was frozen using dry ice and stored at -80 °C. Mice were not perfused prior to brain collection. Frozen hemibrains were homogenized in DPBS using CK14 soft tissue homogenization tubes (Bertin Technologies #P000912-LYSK0-A) to generate 10% (w/v) brain homogenates. For the preparation of detergent-extracted samples, 9 volumes of 10% brain homogenate were mixed with 1 volume of 10X detergent extraction buffer [5% (w/v) sodium deoxycholate and 5% (v/v) NP-40 prepared in DPBS). The samples were incubated on ice for 30 min and then centrifuged at 5,000x *g* for 10 min at 4°C to remove insoluble debris. The supernatant (brain lysate) was collected, and protein concentration was determined using the BCA assay.

### Detergent insolubility assays

For analysis of detergent-insoluble PrP, 50-200 µg of cell or brain lysate was adjusted to a final volume of 50-200 µL of lysis buffer (1X detergent extraction buffer). Unless otherwise indicated, 100 µg of lysate was used for the analysis of detergent-insoluble PrP in all immunoblot samples. The samples were subjected to ultracentrifugation at 100,000x *g* for 1 h at 4 °C. Following ultracentrifugation, the supernatant was removed by gentle aspiration, and the pellet was resuspended in 20 µL of non-reducing 1X Bolt LDS sample buffer (without β-mercaptoethanol). The samples were boiled for 10 min at 95 °C, and then detergent-insoluble protein was loaded onto Bolt Bis-Tris gels and analyzed by immunoblotting.

### Thermolysin digestions

For analysis of thermolysin-resistant PrP species (PrP^AM^), 500 µg of cell or brain lysate was adjusted to a final volume of 200 µL of lysis buffer containing 5-50 µg/mL Thermolysin (TL) (MilliporeSigma #T7902). Unless otherwise indicated, TL was used at a concentration of 20 µg/mL for a final TL-to-protein ratio of 1:125. Samples were digested for 1 h at 37 °C with shaking at 600 rpm, and the digestions were then terminated by the addition of EDTA to a concentration of 5 mM. The samples were then subjected to ultracentrifugation at 100,000x *g* for 1 h at 4 °C. Following ultracentrifugation, the supernatant was removed by gentle aspiration, and the pellet was resuspended in non-reducing 1X Bolt LDS sample buffer (without β-mercaptoethanol). The samples were boiled for 10 min at 95 °C and then stored at -80 °C. Unless otherwise indicated, 200 µg of TL-digested protein was loaded onto Bolt Bis-Tris gels and analyzed by immunoblotting.

### Protein turnover assays

Polyclonal and monoclonal lines of CAD5-PrP^-/-^ cells stably transfected with either WT, D178N-mutant, D178N/G127V-mutant, or E200K-mutant BVPrP(I109) were plated in 6-cm dishes. Cells were cultured for ∼60 h until they reached 90% confluency, and the medium was then replaced with growth medium containing 30 µg/mL cycloheximide (CHX; Sigma #C4859). Cells were lysed with lysis buffer containing protease inhibitor cocktail (Sigma #11873580001) at the indicated time points (0, 2, 4, 6, and 8 h post-CHX treatment). Residual PrP levels were analyzed by immunoblotting. The half-lives of WT and mutant BVPrP(I109) following CHX treatment were determined by fitting residual PrP levels to a one-phase decay model with a fixed plateau of 0 using GraphPad Prism software (version 11.0.2).

### Prion strains

For all *de novo* prion infection experiments, homogenate from persistently infected cultured cells was used as the source of prions (63). Mouse prion strains RML and 22L were respectively derived from prion-infected CAD5 and N2a neuroblastoma cells. The hamster prion strains 263K and HY were propagated in CAD5-PrP^-/-^ cells expressing hamster PrP. The BVPrP-adapted prion strains BV.RML, BV.22L, BV.263K, and BV.HY were derived from prion-infected monoclonal CAD5-PrP^-/-^ cell lines expressing WT BVPrP(I109) (74). To generate prion-infected cellular homogenates, prion-infected cells were cultured in 10-cm tissue culture plates. After reaching confluence, the cells were washed in ice-cold DPBS and then scraped into a small volume of DPBS and collected into CK14 soft tissue homogenization tubes supplemented with additional 0.5 mm zirconia beads (BioSpec #11079105Z). The cells were homogenized using the Minilys apparatus (Bertin Technologies) using 3 cycles of 60 s at maximum speed, with a 10 min incubation on ice between each cycle. Following homogenization, cellular homogenates were aliquoted and stored at -80 °C. Total protein concentrations in cell homogenates were quantified using the BCA assay.

### Cellular prion infections

For *de novo* prion infection experiments, polyclonal lines of stably transfected CAD5-PrP^-/-^ cells were cultured in growth medium which consists of Opti-MEM medium containing 6.5% (v/v) fetal bovine serum, 2 mM GlutaMAX, and 0.2 mg/mL G418 (63). Monoclonal cell lines were infected in the absence of G418. Polyclonal and monoclonal lines of stably transfected CAD5-PrP^-/-^ cells were plated at a density of 4.0-5.0 x 10^4^ cells/well in 24-well plates. The following day, they were cultured in 500 µL of growth media with 100 µg of prion-containing cellular homogenate for 72 h. After 72 h, the cells were washed with DPBS and then passaged in 12-well plates, and they were continuously passaged twice every 3 to 4 days in the 12-well plates and scaled up to 6-cm dishes at the fourth passage. At the fifth passage, the cells were scaled up to 10-cm dishes prior to lysis for the analysis of PK-resistant PrP.

### Proteinase K digestions

For analysis of proteinase K (PK)-resistant PrP, 1 mg of cell lysate was adjusted to a final volume of 400 µL of lysis buffer. Unless otherwise indicated, PK (ThermoFisher #EO0491) was added to a final concentration 50 µg/mL, which corresponds to a PK-to-protein ratio of 1:50. Samples were digested for 1 h at 37 °C with shaking at 600 rpm, and the digestions were then terminated by the addition of PMSF to a concentration of 2 mM. Sarkosyl was added to a final concentration of 2% (v/v), and the samples were then subjected to ultracentrifugation at 100,000x *g* for 1 h at 4 °C. Following ultracentrifugation, the supernatant was removed by gentle aspiration, and the pellet was resuspended in non-reducing 1X Bolt LDS sample buffer (without β-mercaptoethanol). The samples were boiled for 10 min at 95 °C and then stored at -80 °C. PK-digested protein was loaded onto Bolt Bis-Tris gels and analyzed by immunoblotting.

### PNGase F digestions

For analysis of PrP glycosylation in the detergent-insoluble fraction, 100 μg of detergent-insoluble protein from CAD5-PrP^-/-^ cells stably transfected with either D178N-mutant or D178N/G127V-mutant BVPrP(I109) was resuspended in 10 μL of 1x Glycoprotein Denaturing Buffer. The mixture was denatured for 10 min at 95 °C, and then samples were mixed with 2 μL of GlycoBuffer 2 (10x), 2 μL of 10% NP-40, 6 μL of dH_2_O and 1 μL of PNGase F (New England BioLabs #P0708S). As a negative control, 1 µL of dH_2_O was added instead of PNGase F. The mixture was incubated for 1 h at 37 °C, and then the samples were diluted as required into 1X Bolt LDS sample buffer. Deglycosylated samples were analyzed by immunoblotting.

### Treatment of cells with anti-prion compounds

Prion-infected or uninfected CAD5-PrP^-/-^ cells stably expressing WT or mutant BVPrP(I109) were plated at a density of 4.0 x 10^5^ cells in 6-cm dishes. The following day, cells were treated with 2 µM IND24, 2 µM Anle138b, or both compounds (2 µM each) for 72 h. As a control, cells were treated with DMSO. After 72 h post-treatment, the cells were lysed for analysis by immunoblotting.

### Cellular prion seeding activity assay

To generate cellular homogenates containing PrP^AM^, CAD5-PrP^-/-^ cells stably transfected with either WT, D178N-mutant, or D178N/G127V-mutant BVPrP(I109) were cultured in 10-cm dishes until reaching confluence, and then cellular homogenates were aliquoted and stored at -80 °C as described above. Polyclonal lines of CAD5-PrP^-/-^ cells expressing either WT or D178N-mutant BVPrP(I109) were plated at a density of 5.0 x 10^4^ cells/well in 24-well plates. The following day, the cells were cultured in 500 µL of growth media with 100 µg of cellular homogenate containing PrP^AM^ for 72 h. After 72 h, the cells were washed with DPBS and then passaged in 12-well plates. Cells were sequentially expanded from 12-well plates to 6-cm dishes, and then to 10-cm dishes, over two consecutive passages. At the third passage, the cells in 10-cm dishes were lysed for analysis by immunoblotting.

### RT-QuIC assay

Recombinant BVPrP (residues 23-231, M109 isoform) was expressed and purified as described previously (74). Recombinant protein was dialyzed into 10 mM sodium phosphate buffer, pH 7.3, overnight at 4 °C and then ultracentrifuged at 100,000x *g* at 4 °C for 1 h to remove any preformed aggregates. Cell homogenate seeds were prepared from CAD5-PrP^-/-^ cells stably transfected with either D178N-mutant, or D178N/G127V-mutant BVPrP(I109) in 1X PBS with 0.05% (w/v) SDS and 1X N2 supplement (ThermoFisher #17502048) to a concentration of 5.0 ng/μL. Cell homogenates from CAD5-PrP^-/-^ cells or BV.RML-infected CAD5-PrP^-/-^ BVPrP(I109) cells were used as negative or positive controls. The RT-QuIC reaction mixture consisted of 0.1 mg/mL recombinant BVPrP(M109); 10 mM sodium phosphate buffer, pH 7.4; 300 mM NaCl; 1 mM EDTA; and 10 μM thioflavin T. Reactions were carried out in 5 replicates in black, 96-well clear-bottom plates (ThermoFisher #265301). Each well contained 98 μL of reaction mixture and 2 μL of diluted seed. The sealed plates were incubated at 42 °C in a BMG CLARIOstar microplate reader with cycles of 1-minute shake (700 rpm double orbital) and 1-minute rest/read for ∼4 days. The fluorescence excitation and emission wavelengths were 444 ± 5 nm and 485 ± 5 nm, respectively, with a gain setting of 1,600. Lag phases were calculated as previously described (109).

### Immunofluorescence microscopy

CAD5 cells, CAD5-PrP^-/-^ cells, and stably transfected CAD5-PrP^-/-^ lines were plated at density of 1.0 x 10^5^ cells/well in 24-well dishes with a #1.5 glass-like polymer coverslip bottom (Cellvis #P24-1.5P) coated with poly-D-lysine (ThermoFisher #A3890401). Cells were cultured for 48 h until they reached 80-90% confluency. Cells were fixed with 4% (v/v) paraformaldehyde for 15 min and then washed twice with DPBS. Permeabilization was not performed. After washing with DPBS, the cells were blocked in 3% bovine serum albumin (diluted in PBS) for 1 h at room temperature. Cells were then incubated with the anti-PrP antibody POM1 (1:500 dilution) overnight at 4 °C in DPBS containing 3% bovine serum albumin. The following day, the cells were washed twice with DPBS and then incubated with Alexa Fluor 488-conjugated secondary antibody (ThermoFisher #A-11029, 1:500 dilution in DPBS containing 3% bovine serum albumin) for 2 h at room temperature. Subsequently, cells were washed 2 additional times with DPBS. After the final wash, the cells were incubated with DAPI (1 µg/mL in DPBS containing 3% bovine serum albumin) for 10 min and then washed with DPBS. The cells were stored in DPBS and then imaged using a Zeiss LSM880 confocal microscope.

## Statistical analysis

All data was assumed to be normally distributed. When comparing the means of two samples, two-tailed unpaired t tests were used. When comparing the means of three or more samples, one-way ANOVA followed by Dunnett’s) multiple comparisons test was used. All statistical analysis was performed using GraphPad Prism software (version 11.0.2) with a significance threshold of *P* < 0.05.

## Acknowledgements

JCW acknowledges research support from the Canadian Foundation for Innovation/Ontario Research Fund. This research was undertaken, in part, thanks to funding to JCW from the Canada Research Chairs Program.

## Declarations

### Availability of data and material

All data generated or analyzed during this study are included in this published article.

### Author contributions

The study was conceived and designed by GA, GSU, and JCW. GA performed most of the experiments and conducted most of the data analysis. SM and ES collected the brain tissue from the knock-in mice. HA assisted with the RT-QuIC assays. MECB generated the D178N- and E200K-mutant BVPrP(I109) plasmids. SS provided the anti-prion compounds. The first draft of the manuscript was written by GA and JCW, and all authors commented on previous versions of the manuscript. All authors read and approved the final manuscript.

---

**Supplementary Figure S1.**
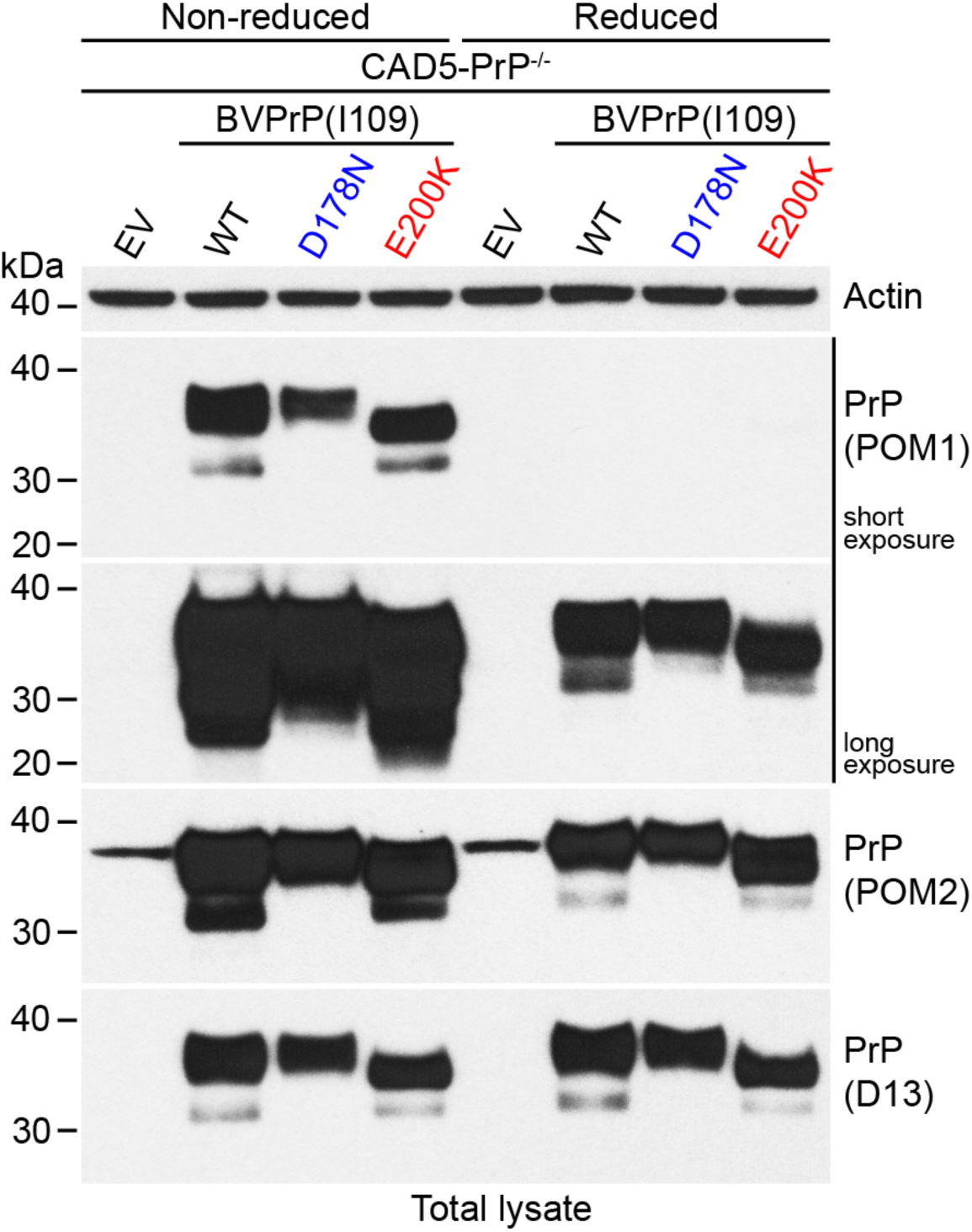
Detection of BVPrP under reducing and non-reducing conditions in polyclonal lines of stably transfected CAD5-PrP^-/-^ cells. Representative immunoblots of PrP in cell lysates from CAD5-PrP^-/-^ cells stably transfected with either empty vector (EV) or with the indicated BVPrP(I109) constructs. In reduced conditions, lysates were treated with β-mercaptoethanol and boiled. The blots were probed with the anti-PrP antibodies POM1, POM2, or D13. The POM1 blot was reprobed with an antibody against actin.

**Supplementary Figure S2.**
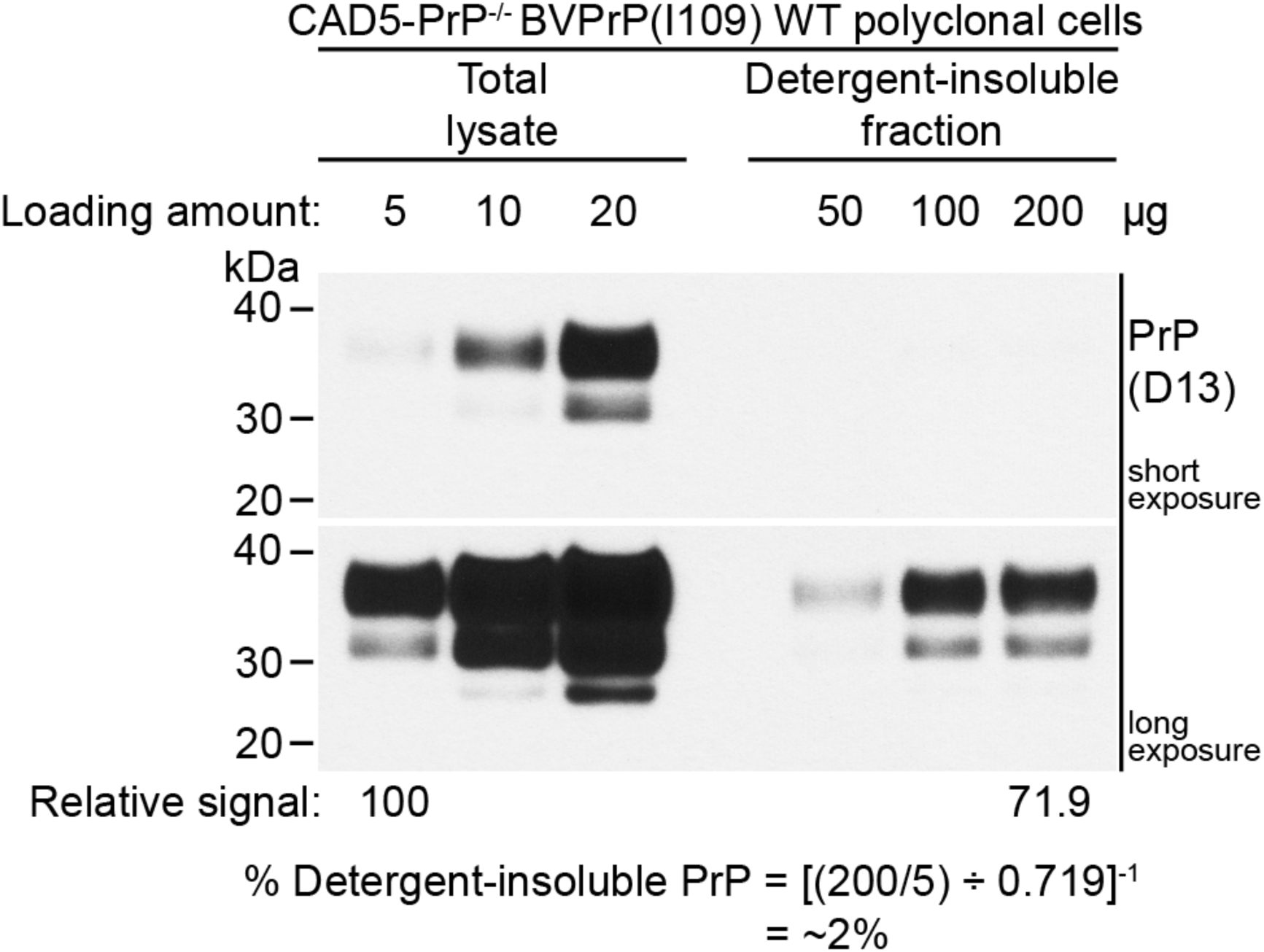
Low levels of detergent-insoluble PrP species are present in cells expressing WT BVPrP(I109). Immunoblot of total and detergent-insoluble PrP species in cell lysates from polyclonal CAD5-PrP^-/-^ cells stably transfected WT BVPrP(I109). By loading different amounts of the total and detergent-insoluble fractions, the proportion of WT BVPrP(I109) species that are detergent-insoluble was calculated to be ∼2%. PrP was detected using the antibody D13.

**Supplementary Figure S3.**
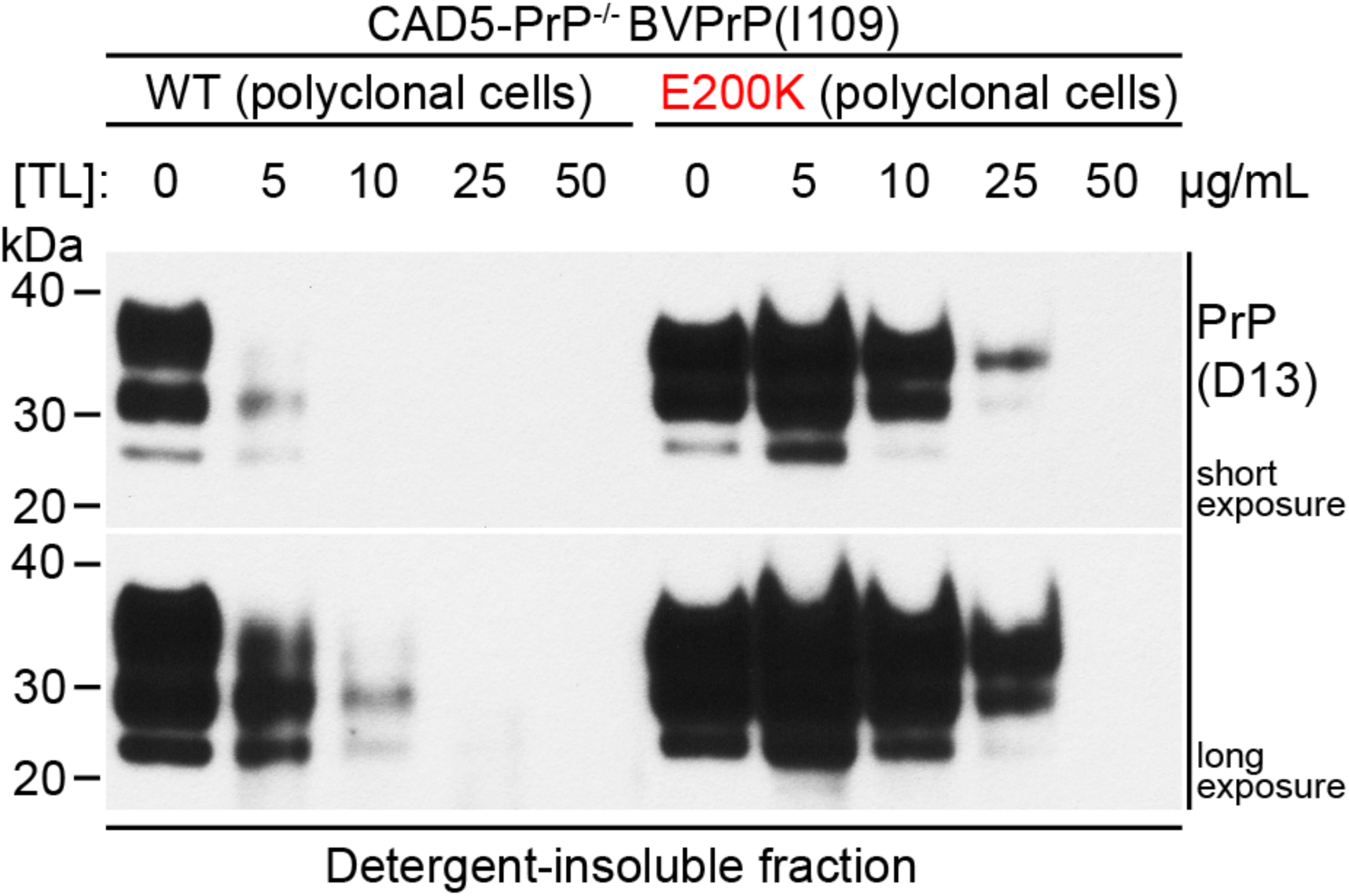
Thermolysin sensitivity of PrP species in cells stably expressing WT or E200K-mutant bank vole PrP. Representative immunoblot of detergent-insoluble PrP species in cell lysates from CAD5-PrP^-/-^ cells stably transfected with either WT or E200K-mutant BVPrP(I109) following treatment with the indicated concentrations of thermolysin (TL). TL-resistant PrP was detected using the antibody D13.

**Supplementary Figure S4.**
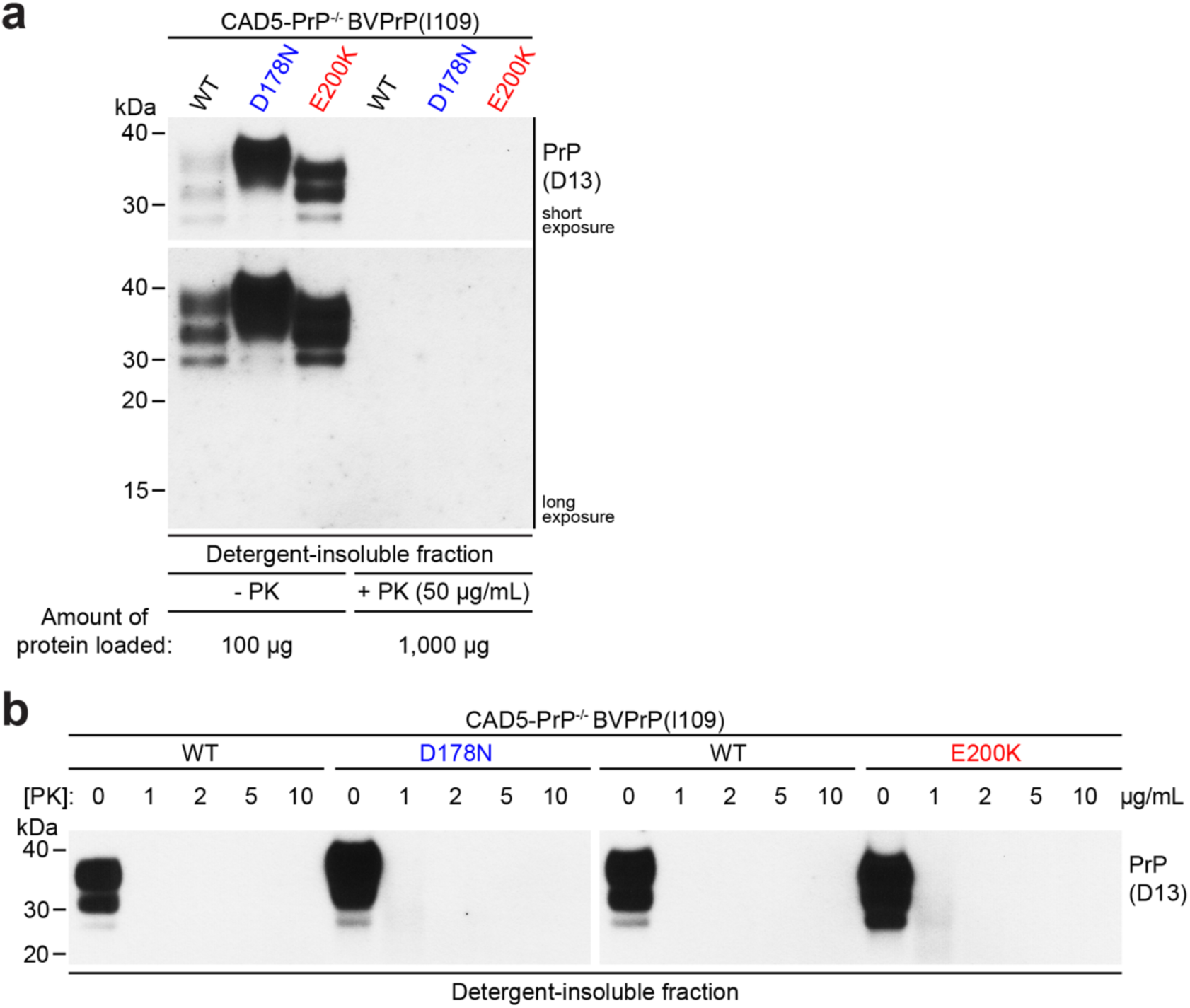
Polyclonal lines of stably transfected CAD5-PrP^-/-^ cells expressing mutant bank vole PrP do not exhibit proteinase K-resistant PrP. **a**) Representative immunoblot of detergent-insoluble PrP species in lysates from stably transfected CAD5-PrP^-/-^ cells that were either treated (+ PK) or not treated (- PK) with 50 µg/mL proteinase K. **b**) Representative immunoblots of detergent-insoluble PrP species in cell lysates from CAD5-PrP^-/-^ cells stably transfected with either WT, D178N-mutant, or E200K-mutant BVPrP(I109) following treatment with the indicated concentrations of proteinase K. In both panels, PrP was detected using the antibody D13.

**Supplementary Figure S5.**
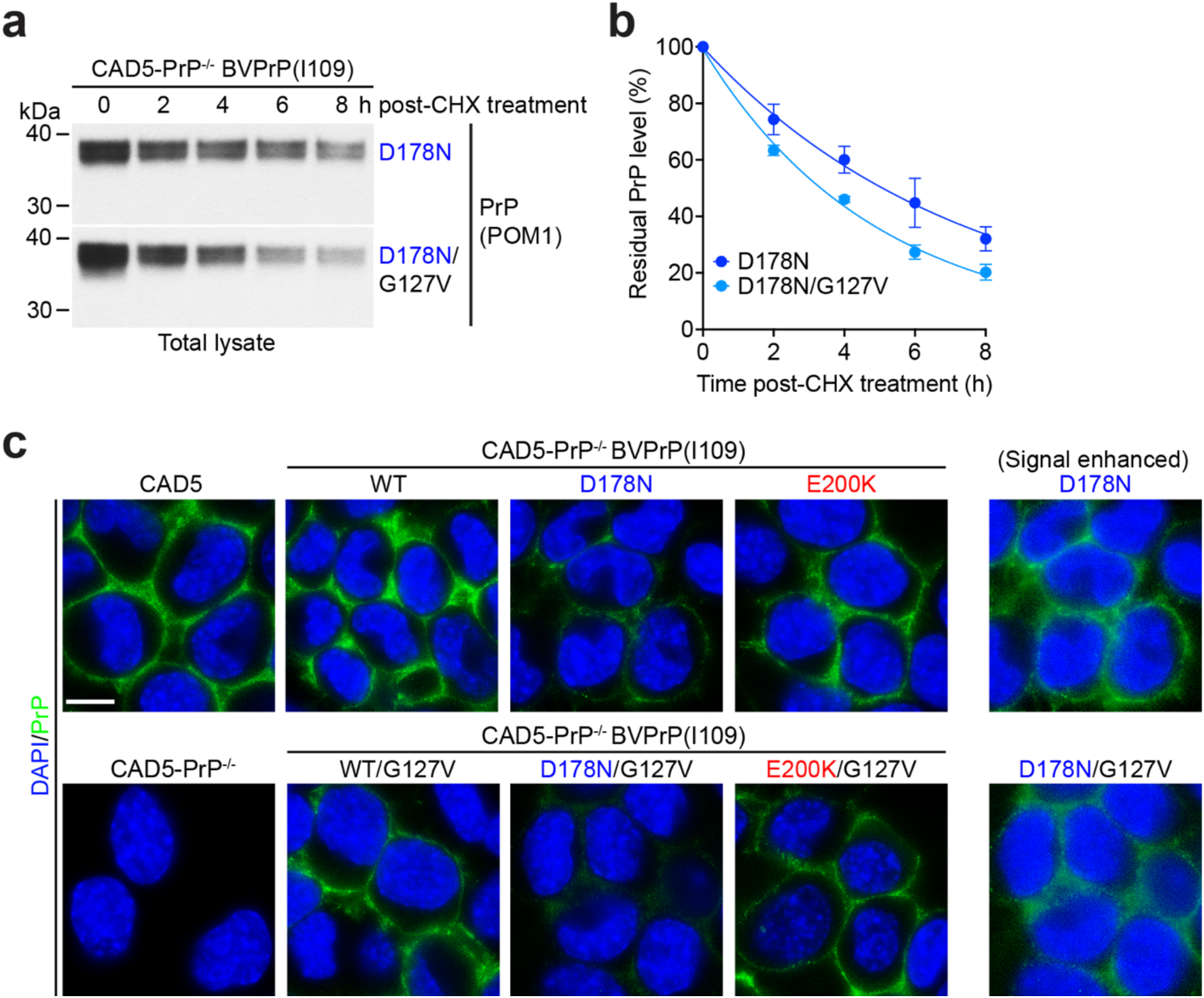
Stability of D178N-mutant BVPrP(I109) containing G127 or V127 and cell surface localization of BVPrP(I109) variants. **a**) Immunoblots of residual PrP levels in lysates from stably transfected polyclonal CAD5-PrP^-/-^ lines expressing D178-mutant BVPrP(I109) with or without the G127V mutation treated with cycloheximide (CHX) for the indicated amounts of time. **b**) Quantification of residual PrP levels in the polyclonal cell lines following treatment with CHX for the indicated amounts of time (n = 3 independent experiments). The half-life of D178N-mutant BVPrP(I109) was calculated to be 5.1 h whereas the half-life of D178N-mutant BVPrP(I109) containing G127V was calculated to be 3.4 h. **c**) Immunofluorescence images of polyclonal lines of CAD5-PrP^-/-^ cells stably expressing the indicated BVPrP(I109) variants. CAD5 and CAD5-PrP^-/-^ cells are included as controls. All images were processed identically, except for the rightmost column in which the signal was enhanced to reveal cell-surface PrP staining in cells expressing D178N- or D178N/G127V-mutant BVPrP(I109). PrP expression (green) was revealed using the anti-PrP antibody POM1 and nuclei were stained with DAPI (blue). Scale bar = 10 μm (applies to all images).

**Supplementary Figure S6.**
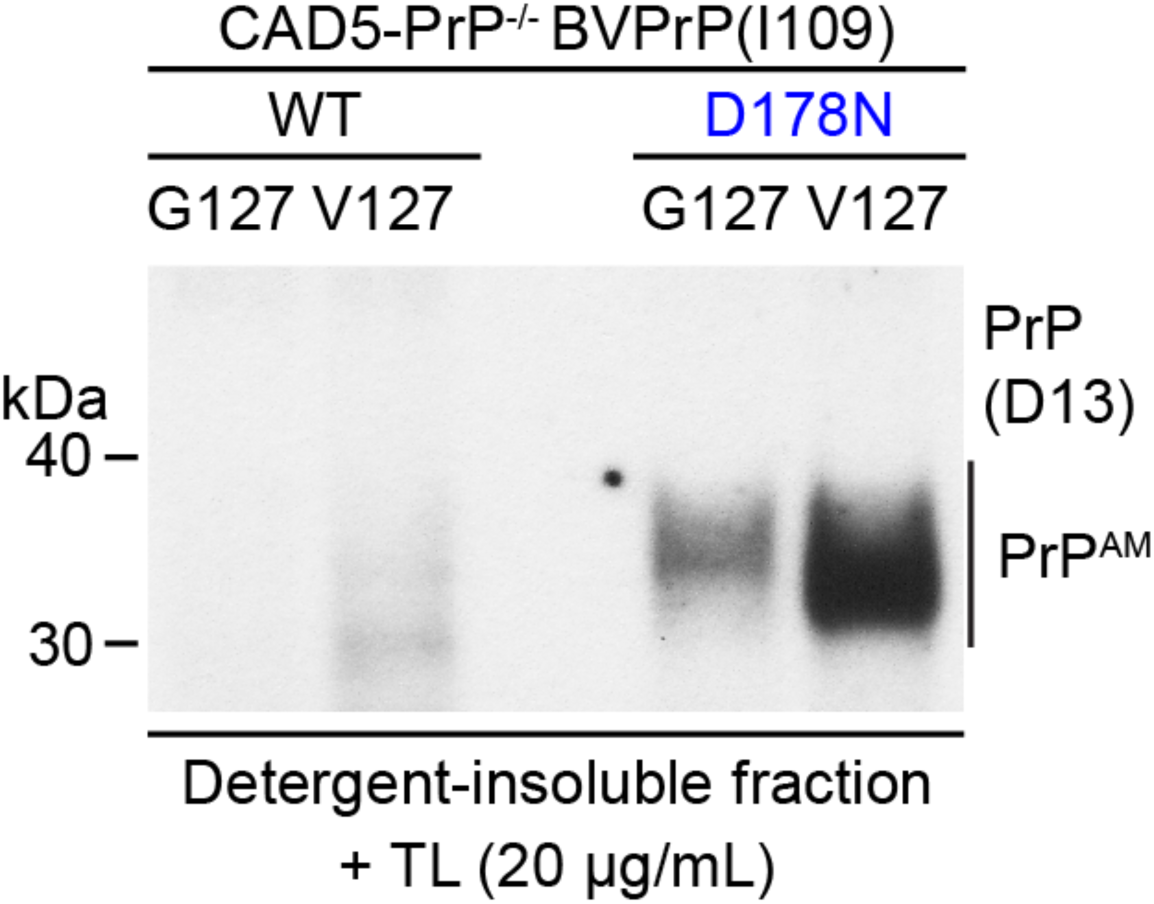
The G127V substitution also induces formation of PrP^AM^ when present in wild-type bank vole PrP. Representative immunoblot of detergent-insoluble, TL-resistant PrP species in cell lysates from CAD5-PrP^-/-^ cells stably transfected with either WT or D178N-mutant BVPrP(I109), each containing either glycine or valine at codon 127. TL-resistant PrP was detected using the antibody D13.

**Supplementary Figure S7.**
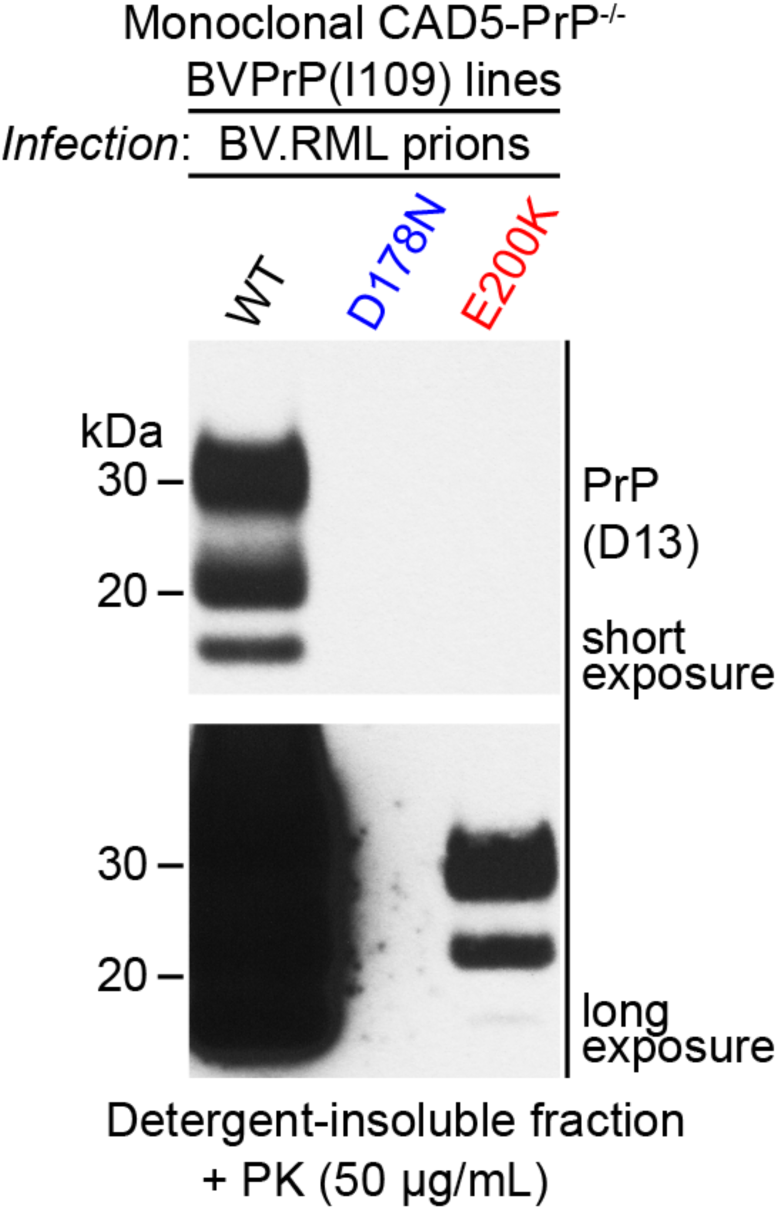
Low amounts of PK-resistant PrP in monoclonal CAD5-PrP^-/-^ cells expressing E200K-mutant BVPrP(I109) following challenge with BV.RML prions. Representative immunoblot of PK-resistant PrP levels in lysates from monoclonal CAD5-PrP^-/-^ cell lines expressing either WT, D178N-mutant, or E200K-mutant BVPrP(I109) after 5 passages following infection with BV.RML prions. The two images represent different exposures of the same blot.

